# Mitigation of chromosome loss in clinical CRISPR-Cas9-engineered T cells

**DOI:** 10.1101/2023.03.22.533709

**Authors:** Connor A. Tsuchida, Nadav Brandes, Raymund Bueno, Marena Trinidad, Thomas Mazumder, Bingfei Yu, Byungjin Hwang, Christopher Chang, Jamin Liu, Yang Sun, Caitlin R. Hopkins, Kevin R. Parker, Yanyan Qi, Ansuman T. Satpathy, Edward A. Stadtmauer, Jamie H.D. Cate, Justin Eyquem, Joseph A. Fraietta, Carl H. June, Howard Y. Chang, Chun Jimmie Ye, Jennifer A. Doudna

## Abstract

CRISPR-Cas9 genome editing has enabled advanced T cell therapies, but occasional loss of the targeted chromosome remains a safety concern. To investigate whether Cas9-induced chromosome loss is a universal phenomenon and evaluate its clinical significance, we conducted a systematic analysis in primary human T cells. Arrayed and pooled CRISPR screens revealed that chromosome loss was generalizable across the genome and resulted in partial and entire loss of the chromosome, including in pre-clinical chimeric antigen receptor T cells. T cells with chromosome loss persisted for weeks in culture, implying the potential to interfere with clinical use. A modified cell manufacturing process, employed in our first-in-human clinical trial of Cas9-engineered T cells,^1^ dramatically reduced chromosome loss while largely preserving genome editing efficacy. Expression of p53 correlated with protection from chromosome loss observed in this protocol, suggesting both a mechanism and strategy for T cell engineering that mitigates this genotoxicity in the clinic.

## Introduction

The precision of CRISPR-Cas9 genome editing is paramount to ensure clinical safety and avoid unintended and permanent genotoxicities. Promiscuous off-target genome editing at unintended sites has been extensively studied^2–4^ and mitigated^5, 6^ *in vitro* and *in vivo*. However, unintended chromosomal abnormalities following on-target genome editing, such as chromosome loss, have not been systematically investigated or prevented. Thus, these potential concerns for genome editing continue to persist, including in the clinic, at an unknown frequency.

T cells have been extensively engineered using Cas9 to develop potent immunotherapies for cancer^1, 7–9^ and autoimmune diseases.^10, 11^ In a previous study, low-level chromosome 14 aneuploidy was detected in primary human T cells after Cas9-mediated genome editing of the *T cell receptor alpha constant* (*TRAC*) gene using one clinically relevant guide RNA (gRNA).^12^ However, the extent to which chromosome loss occurs at other target sites, the determinants of this phenomenon, the behavior of T cells with Cas9-induced chromosome loss, and the clinical relevance of these findings remain unknown. Along with understanding this phenomenon, strategies to reduce or eliminate chromosome loss as an outcome of genome editing would improve the safety of engineered T cell therapies for patients.

Here, we analyzed chromosome loss following Cas9-induced genome editing at hundreds of sites across every somatic chromosome in the human genome to determine the frequency, determinants, and consequences of this phenomenon. We found Cas9-induced chromosome loss was a generalizable phenomenon, although it was specific to the chromosome targeted by Cas9, and was prevalent at sites across the genome. T cells with Cas9-induced chromosome loss had a fitness and proliferative disadvantage, yet could persist over multiple weeks of *ex vivo* culture. Surprisingly, chromosome loss also occurred during pre-clinical production of chimeric antigen receptor (CAR) T cells but was minimal or undetectable in Cas9-edited patient T cells from a first-in-human phase 1 clinical trial. Further experimentation demonstrated that a modified T cell editing protocol employed in our clinical trial increased levels of the DNA damage response protein p53 while decreasing chromosome loss, suggesting a possible mechanism for Cas9-induced chromosome loss and an unexpected strategy to avoid this unintended outcome in patients.

## Results

### Single-cell RNA sequencing reveals chromosome loss in Cas9-edited T cells

The *TRAC* locus, which encodes the T cell receptor (TCR) responsible for CD4 and CD8 T cell reactivity to peptide-MHC complexes, is of immense interest for genome editing applications in human T cells. For engineered adoptive T cell therapies, where T cells are armed with a transgenic receptor for targeted immunotherapy, disrupting endogenous TCR expression limits graft-versus-host toxicity associated with mispairing of transgenic and endogenous TCRs.^13^ Additionally, abrogating the TCR is an important step towards developing allogeneic “off-the-shelf” T cell therapies that could reduce cell manufacturing costs and expand patient accessibility.^14^

To quantify chromosome loss after genome editing of *TRAC*, primary human T cells were nucleofected with *S. pyogenes* Cas9 ribonucleoprotein (RNP) including one of 11 different gRNAs tiled across the first exon of *TRAC* or a non-targeting gRNA (Fig. 1A). Reduced TCR expression occurred in 60-99% of all cells as measured by flow cytometry (Fig. S1A) and editing at the *TRAC* locus was observed in 62-97% of all genomic DNA sequencing reads (Fig. S1B), depending on the gRNA. Four days after Cas9 RNP introduction, T cells were subjected to multiplexed single-cell RNA sequencing (scRNA-seq) to detect reduced transcript levels resulting from chromosome loss caused by Cas9-mediated double-strand DNA breaks (DSBs) (Fig. S1C).^15^ Transcriptome-wide analysis using existing computational methods revealed a reduction in gene dosage, specifically on chromosome 14, in cells with a *TRAC*-targeting gRNA compared to cells with a non-targeting gRNA (Fig. 1B).^16^ We further estimated the distribution of breakpoint loci across chromosome 14, finding the highest frequency to be near the known genomic location of our *TRAC*-targeting gRNAs (Fig. 1C). We observed cells with lower gene dosage originating at the Cas9 target site (partial chromosome loss) as well as cells with lower gene dosage across the entire chromosome (whole chromosome loss) (Fig. 1D). Overall, ∼5-20% of T cells exhibited partial or whole loss of chromosome 14 depending on the *TRAC*-targeting gRNA (Fig. 1E, S1D, S1E, and S1F).

**Figure 1:**
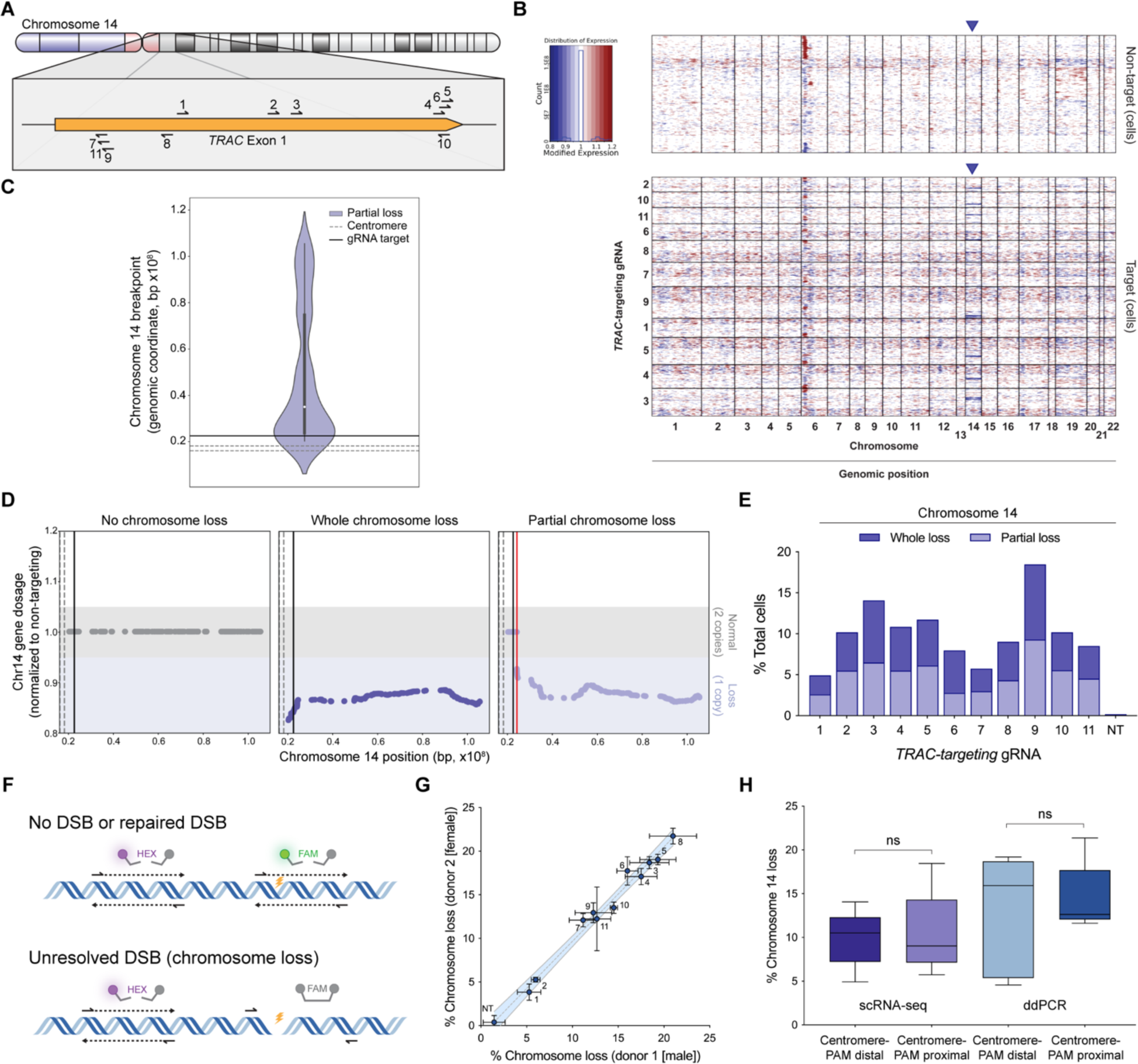
CRISPR-Cas9 genome editing of *TRAC* results in whole and partial chromosome loss. **(A)** Cas9 gRNA target sites tiled across the first exon of *TRAC* on chromosome 14. **(B)** Gene dosage from transcriptome-wide scRNA-seq of T cells treated with Cas9 and a non-targeting gRNA (top heatmap) or *TRAC*-targeting gRNA (bottom heatmap). Each individual row corresponds to a single cell and each column corresponds to a specific gene and its genomic position, grouped by chromosome (outlined in black). Red represents increase in gene dosage while blue represents decrease in gene dosage. Rows outlined in black represent cells treated with different *TRAC*-targeting gRNAs. Blue arrows highlight chromosome 14, where *TRAC* is located. **(C)** Distribution of computationally predicted chromosome 14 breakpoints in cells predicted to have a chromosomal loss event. The distribution is an aggregate of 11 different *TRAC*-targeting gRNAs (all within ∼300 bp) in cells with partial chromosome loss. **(D)** Representative single cell chromosome 14 gene dosage plots illustrating a cell with no chromosome loss (left), whole chromosome loss (middle), or partial chromosome loss (right). Gene dosage was normalized to non-targeting samples. Gray shaded area (gene dosage of 0.95-1.05) represents normal gene dosage (2 copies). Blue shaded area (gene dosage of <0.95) represents reduction in gene dosage (1 copy). Dotted lines represent the centromere, black lines represent the Cas9 target site, and the red line represents the computationally predicted breakpoint. **(E)** Quantification of whole and partial chromosome 14 loss across all gRNAs from scRNA-seq. NT indicates non-targeting gRNA. **(F)** Schematic of ddPCR assay to measure chromosome loss. The yellow lightning bolt represents the Cas9 target site. The detection of both HEX and FAM probes indicates no DSB or repaired DSB (top illustration). The detection of the HEX probe but not the FAM probe indicates an unresolved DSB that represents chromosome loss (bottom illustration). **(G)** Quantification of chromosome loss at the Cas9 target site across all gRNAs from ddPCR (n = 3, n = 2 biological donors). Numbers next to each point represent the *TRAC*-targeting gRNA. NT indicates non-targeting gRNA and represents samples from four different ddPCR amplicons. Error bars represent the standard deviation from the mean. Dashed line represents linear regression line of best fit and shaded region represent 95% confidence intervals (Slope = 1.082, R^2^ = 0.9853). **(H)** Comparison of chromosome 14 loss between *TRAC*-targeting conditions where the PAM is distal (Centromere-PAM distal) or proximal (Centromere-PAM proximal) to the centromere relative to the gRNA spacer sequence. Chromosome 14 loss was measured by scRNA-seq (n = 1 biological donor) or ddPCR (n = 3, n = 2 biological donors). *P*-values are from Welch’s unpaired t-tests and are from left to right 0.8689 and 0.7338. ns = not significant.

### DNA-based droplet digital PCR is an orthogonal method to detect chromosome loss

As an orthogonal approach to detect chromosome loss, we used droplet digital PCR (ddPCR) to directly quantify genomic DNA copy number, eliminating potential interference from transcriptional or epigenetic factors that may affect the scRNA-seq results. Two primer/probe sets were designed to amplify nearby regions of the target gene, one as a control (HEX) and the other spanning the Cas9 gRNA target site (FAM) so that a DSB and chromosome loss would inhibit amplicon generation (Fig. 1F). Three days after Cas9 nucleofection, genomic DNA harvested from *TRAC*-targeted T cells resulted on average in ∼4-22% chromosome loss; these losses were highly reproducible across biological T cell donors (Fig 1G). Importantly, primers and probes in the amplicon spanning the intended Cas9 target site were positioned to avoid binding site disruption by small insertions and deletions (indels), the most common outcome after Cas9 genome editing.

Based on the observation that Cas9 preferentially remains bound to the non-protospacer adjacent motif (PAM) side of the target DNA after cleavage,^17, 18^ we wondered whether orientation of the PAM relative to the centromere influenced chromosome loss (Fig. S1G). However, we found no significant difference between targets where the PAM was distal or proximal to the centromere, relative to the gRNA spacer sequence (Fig. 1H). Furthermore, the rates of chromosome 14 loss measured by scRNA-seq or ddPCR did not correlate with the efficacy of TCR disruption or genomic position targeted by different gRNAs (Fig. S1H and S1I).

### Target-specific chromosome loss is a general phenomenon following genome editing

To determine the generality of chromosome loss after genome editing, we conducted a comprehensive CRISPR screen using a library of 384 unique gRNAs targeting 3-7 genes on every somatic chromosome (92 genes in total) with four unique gRNA sequences targeting each gene (Fig. 2A, S2A, and S2B). CROP-seq was used to track gRNAs delivered to individual cells in our experiment because it avoids the template switching observed with other methods^19, 20^ and because it has been successfully deployed in primary human T cells.^21^ Targets relevant to T cell genome engineering were prioritized, such as *TRAC*,^1, 22, 23^ *TRBC*,^1^ *PDCD1*,^1^ *B2M*,^24^ *IL2RA*,^23^ *CXCR4*,^25^ and *CIITA*,^26^ as well as other targets of interest for clinical genome editing such as *BCL11A*^27, 28^ and *HBB*^29, 30^ for the treatment of sickle cell disease, *TTR* to treat transthyretin amyloidosis,^31^ *HTT* to treat Huntington’s disease,^32^ and *SERPINA1* to treat alpha-1-antitrypsin deficiency.^33^

**Figure 2:**
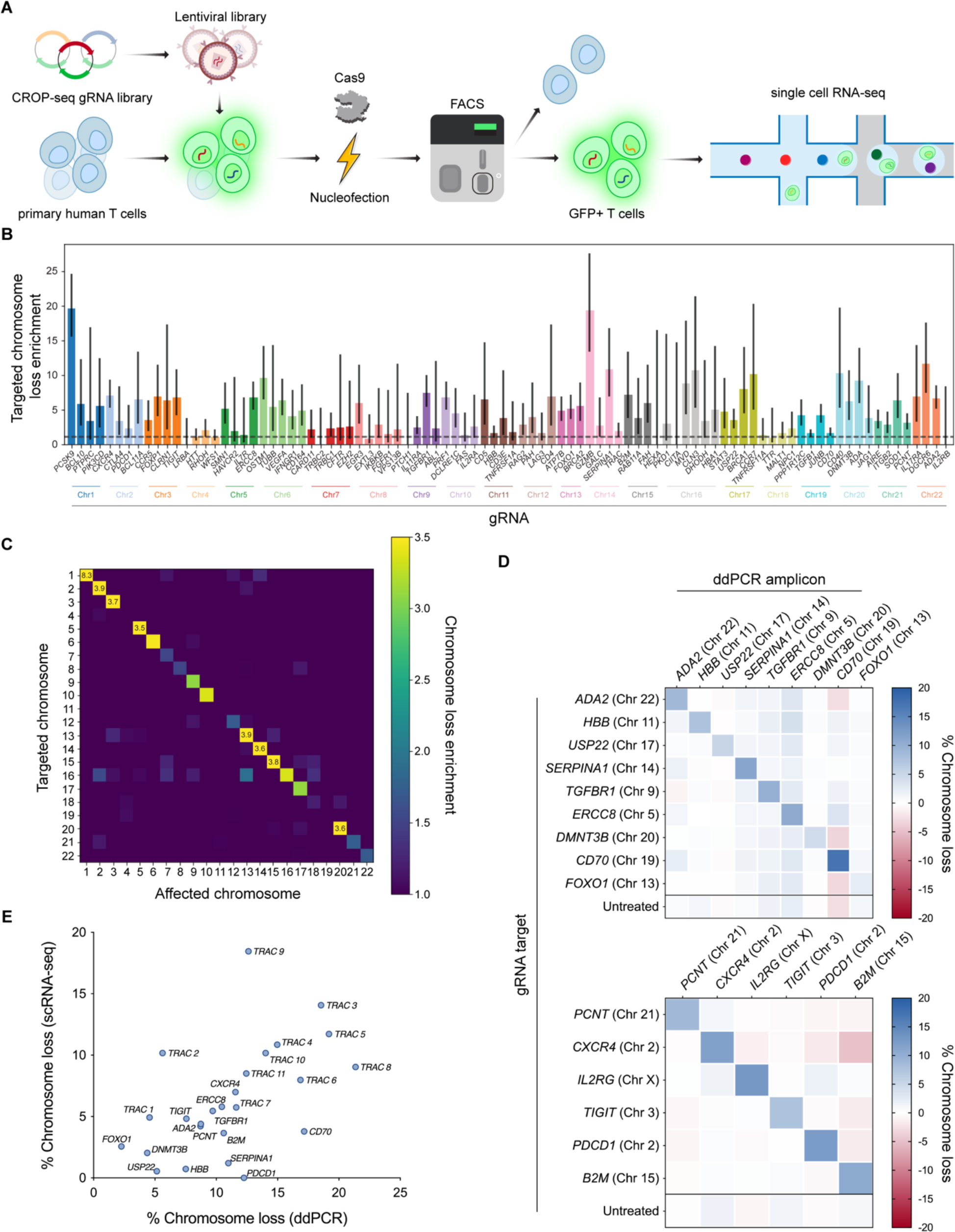
Genome-scale CRISPR-Cas9 screen reveals target-specific chromosome loss. **(A)** Workflow for a CRISPR-Cas9 screen to estimate chromosome loss in T cells. Primary human T cells were transduced with a CROP-seq lentiviral library expressing one of 384 gRNAs. Cells were then nucleofected with Cas9 protein, before GFP+ cells (co-expressed on the CROP-seq gRNA vector) were enriched via fluorescence-activated cell sorting. Enriched cells were subject to scRNA-seq and downstream analysis. **(B)** Quantification of targeted chromosome loss enrichment for each target gene. Each of the 92 bars represents the combination of four unique gRNAs targeting the same gene. Chromosome loss enrichment was calculated relative to the baseline loss per chromosome in cells containing a gRNA targeting a different chromosome. Error bars represent 95% confidence intervals. **(C)** Chromosomal loss enrichment at each somatic chromosome across all gRNAs. Rows represent the chromosome targeted by the Cas9 gRNA. Columns represent the chromosome analyzed for chromosome loss. **(D)** Chromosome loss measured by ddPCR at 15 different Cas9 target sites across the genome. Rows titles indicate the identity of the gRNA used. Column titles indicate the site in the genome that was analyzed via ddPCR. Heatmap values represent the mean of replicates (n = 3, except n = 2 for *B2M* target column). **(E)** Correlation between chromosome loss from 25 gRNAs as measured by scRNA-seq and ddPCR. Spearman’s correlation = 0.59, \*\**P* = 0.0017 (two-tailed).

Using our previously established computational pipeline on the CROP-seq dataset, we determined the breakpoints and gene dosage for 92 different genes targeted by Cas9 (Fig. S1C, S1D, S1E, and S1F). For numerous genes targeted in our screen, we observed significant enrichment for loss of the targeted chromosome in cells with a corresponding gRNA compared to cells with a gRNA targeting a different chromosome (Fig. 2B). Chromosome loss above background levels occurred with 55% (201/364) of all gRNAs, for 89% (82/92) of all genes targeted, and in 100% (22/22) of all chromosomes targeted. Across all gRNAs, 3.25% of targeted cells had detectable whole or partial chromosome loss (Fig. S3A). For cells with a non-targeting gRNA, no detectable chromosome loss was identified on any somatic chromosome. Enrichment for chromosome loss was much higher in chromosomes targeted by Cas9 compared to non-targeted chromosomes (Fig. 2C), suggesting that this phenomenon is an outcome of target-specific cleavage during Cas9-mediated genome editing.

We validated this genome-scale chromosome loss by selecting 15 gRNAs from the library and individually nucleofecting them as Cas9 RNPs into T cells. ddPCR was used to measure chromosome loss at various sites in the genome and showed greater levels of chromosome loss at the targeted site compared to non-targeted sites for nearly all gRNAs (Fig. 2D). Additionally, rates of chromosome loss were highly correlated (Spearman’s correlation = 0.59) with those estimated by scRNA-seq (Fig. 2E).

Further analyses of the CROP-seq screen to investigate the contribution of the Cas9 gRNA sequence revealed no influence by gRNA binding orientation (Fig. S3B), nucleotide sequence (Fig. S3C and S3D), or GC content on chromosome loss (Fig. S3E). The computationally predicted dominant end-joining repair pathway, namely non-homologous end joining (NHEJ) or microhomology-mediated end joining (MMEJ), for each gRNA target also did not show a strong correlation with chromosome loss (Fig. S3F and S3G). However, we did observe a moderate correlation between the distance of each targeted gene from the centromere and the rate of chromosome loss induced (Fig. S3H), with gRNAs targeting closer to the centromere showing higher levels of chromosome loss.

### Chromosome loss accompanies transcriptional signatures of DNA damage response, apoptosis, and quiescence

To assess the functional effects of Cas9-induced chromosome loss, we performed differential gene expression analysis between Cas9-edited T cells with or without chromosome loss. We identified genes that were differentially expressed across groups of cells with different chromosomes lost (Fig. 3A and S4A). *CD70*, for example, was significantly upregulated in every group of cells with chromosome loss regardless of which chromosome was lost, and *MDM2* was significantly upregulated in every group of cells with chromosome loss except those that lost chromosomes 13 or 18. Meanwhile, *PHGDH* was downregulated in every group of cells with chromosome loss except those that lost chromosomes 6, 10, 18, or 21. The numerous genes that were differentially expressed across cells with various chromosomes lost suggests that these findings are not a result of expression changes from lowered dosage of the target gene, but a general influence of Cas9-induced chromosome loss.

**Figure 3:**
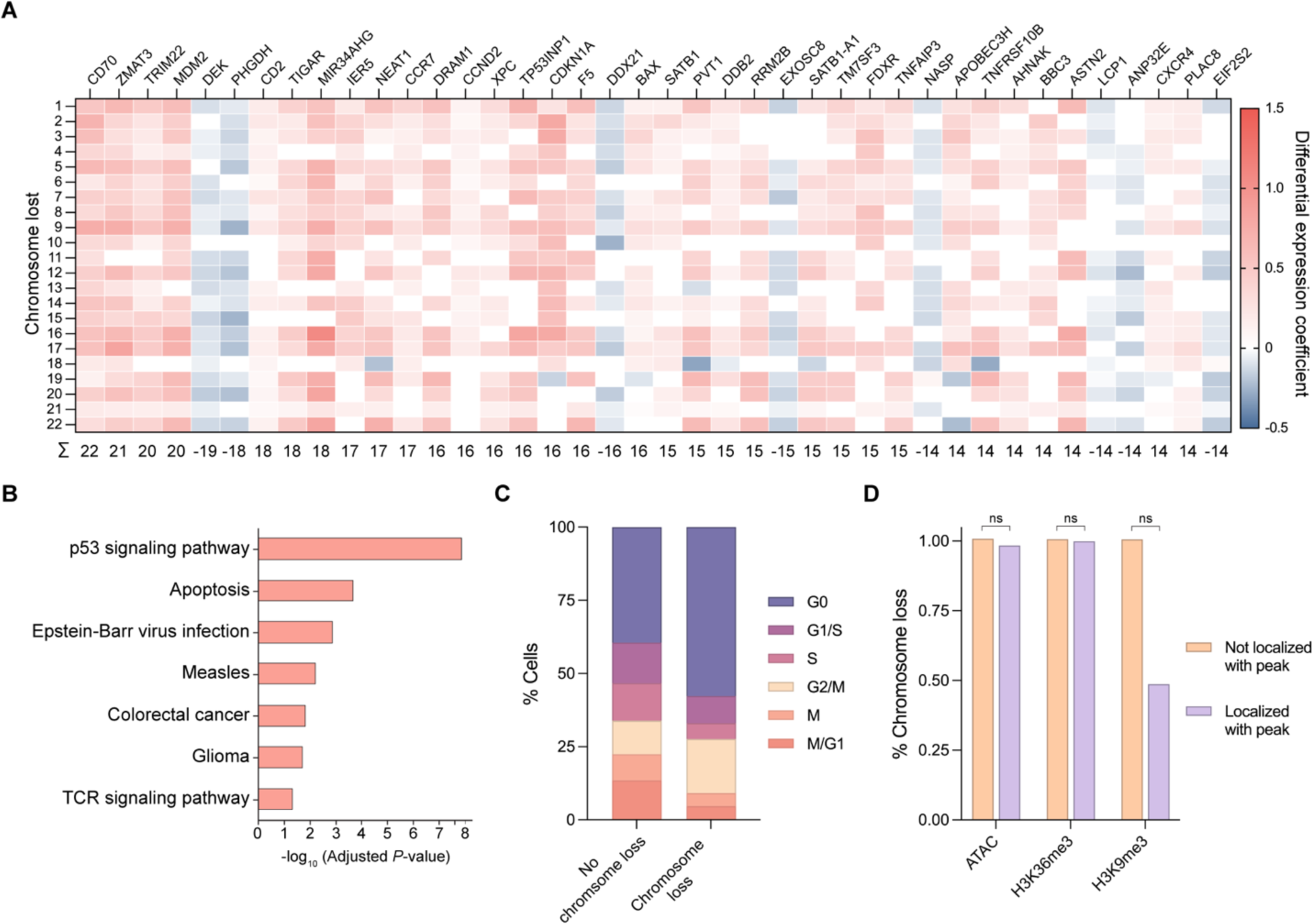
Genetic and epigenetic factors influence Cas9-induced chromosome loss. **(A)** Heatmap of differentially expressed genes in cells with chromosome loss compared to cells without chromosome loss. Cells with chromosome loss were divided into 22 groups depending on which somatic chromosome was lost (rows), and differentially expressed genes were individually investigated (columns). Upregulated genes are shown in red while downregulated genes are shown in blue. Genes were given a score of 1 (upregulated), −1 (downregulated), or 0 (no difference) for each chromosome loss group. Summed gene scores across all chromosome loss groups is shown below; genes with a score >|13| are displayed. **(B)** Gene ontology analysis based on differential gene expression. The most significantly upregulated modules are displayed. **(C)** Cell cycle analysis based on expression profiles. The percentage of cells in each cell cycle phase were quantified for cells with no chromosome loss or cells with chromosome loss. **(D)** Influence of epigenetic marks on chromosome loss. The gRNA sequence for cells with or without chromosome loss was analyzed for localization within ± 75 bp of an epigenetic marker peak. *P*-values were calculated using a two-sided Fisher’s Exact Test and are from left to right (ATAC) 0.365496, (H3K36me3) 0.789824, and (H3K9me3) 0.305706. ns = not significant.

Previous studies have demonstrated that a single Cas9-induced DSB can lead to p53 upregulation.^34^ Consistent with this finding, gene ontology analysis revealed the p53 DNA damage response and general apoptosis pathways were the most significantly overexpressed gene modules in cells with Cas9-induced chromosome loss (Fig. 3B). We also observed an increase in cell cycle markers associated with the G0 phase and a decrease in those associated with the S phase for T cells with chromosome loss compared to those without (Fig. 3C, S4B, and S4C), suggesting p53-induced cell cycle arrest. The results of both the differential gene expression analysis and cell cycle analysis indicate reduced fitness in T cells with Cas9-induced chromosome loss.

We further investigated the relationship between epigenetic markers and chromatin accessibility with Cas9-induced chromosome loss in primary human T cells; however, no significant correlation was found between these epigenetic factors and chromosome loss (Fig. 3D).

### T cells with chromosome loss persist ex vivo but with reduced fitness and proliferation

To determine whether T cells with Cas9-induced chromosome loss persist over time, we used ddPCR to measure the extent of chromosome loss at various timepoints during *ex vivo* culture. We chose timepoints over 2-3 weeks, which is a similar length of *ex vivo* culture compared to the manufacturing protocols of clinical trials with Cas9-edited T cells.^1, 7^ As expected, T cells treated with Cas9 RNP targeting *TRAC* showed the highest levels of DSBs one day after nucleofection (Fig. 4A), when Cas9 RNP is still present within cells and DNA repair is ongoing.^35, 36^ By day three post-treatment, DSBs plateaued until day 14 with most conditions showing a slight downward trend (Fig. 4A). Since Cas9 RNP-mediated cleavage and DNA repair go to completion within 24 hours,^37^ we posited that DSBs measured at day three and beyond are from unrepaired DSBs which we considered chromosome loss.

**Figure 4:**
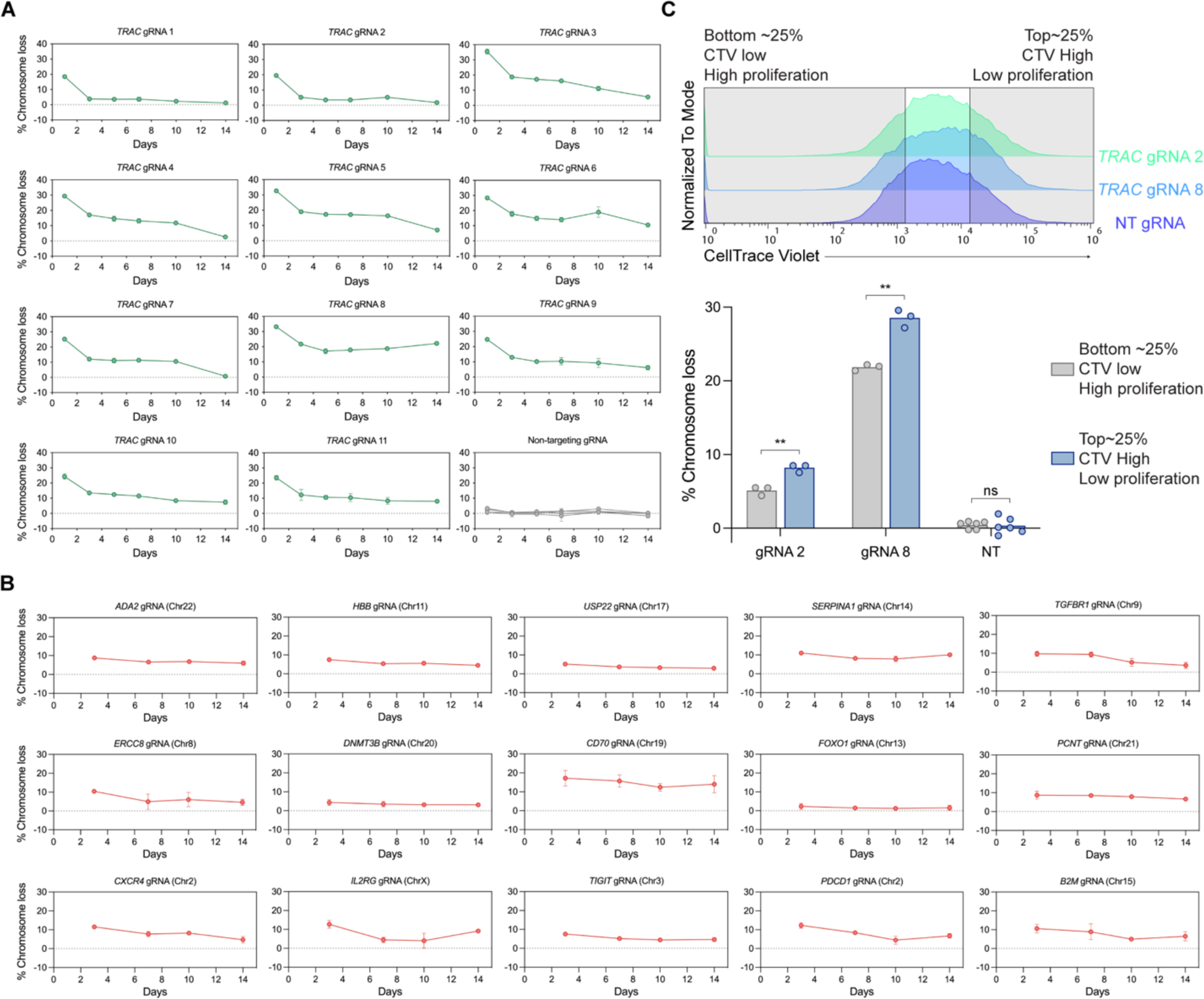
Cas9-induced chromosome loss persists for weeks but results in reduced fitness and proliferation. **(A)** ddPCR measurements of chromosome loss at the Cas9 *TRAC* target site over 14 days. Error bars represent the standard deviation from the mean (n = 3). Day 3 results were additional used as the donor 2 (female) results shown in Fig. 1g. **(B)** ddPCR measurements of chromosome loss for 15 different gRNAs targeted to sites across the genome over 14 days. Error bars represent the standard deviation from the mean (n = 3, except n = 2 for *B2M*). Day 3 results were additionally used for the diagonal values in the heatmaps of Fig. 2d. **(C)** Measurement of chromosome loss across T cells of varying proliferative capacity. T cells were stained with CellTrace Violet (CTV) and cultured for five days before sorting the top and bottom quartile (top panel). ddPCR was used to measure chromosome loss in lowly proliferative (CTV high) and highly proliferative (CTV low) populations (bottom panel). NT = non-targeting gRNA. Non-targeted samples evaluated for chromosome loss at the gRNA 2 or gRNA 8 amplicon were combined into a single column (n = 3 for each of the two different ddPCR amplicons). *P*-values are from Welch’s unpaired t-tests and from left to right are 0.002970, 0.002970, and 0.275572.

We evaluated the temporal dynamics of chromosome loss over a longer period by repeating the experiment over the course of 21 days using four of the 11 *TRAC*-targeting gRNAs. Again, levels of chromosome loss showed a slight reduction over the three weeks, with detectable chromosome loss at the last timepoint remaining above that of non-targeting controls (Fig. S4D).

To test the possibility that targeting a specific gene or chromosome may affect chromosome loss or T cell viability, we repeated the Cas9 nucleofection and genomic DNA ddPCR with 15 gRNAs targeting other genes throughout the genome. Culturing the genome edited T cells for 14 days and measuring chromosome loss at various timepoints throughout, we once again observed a gradual reduction in chromosome loss over time (Fig. 4B). These findings show that chromosome loss in T cells is observable as far out as 2-3 weeks, across multiple targeted genes and chromosomes. However, the gradual decrease in chromosome loss over time suggests a fitness disadvantage for T cells with this genomic aberration.

We additionally tested whether chromosome loss was associated with a proliferative disadvantage in T cells. *TRAC*-edited T cells were stained with a cell proliferation dye and cultured for five days. Cells that underwent the highest and lowest amounts of proliferation, based on dye intensity, were sorted and chromosome loss was measured between the two groups. Chromosome loss in the highest proliferating quartile was identifiable but statistically lower than chromosome loss in the lowest proliferating quartile (Fig. 4C), which suggests that Cas9-induced chromosome loss confers a proliferative disadvantage.

### Gene insertion via homology-directed repair results in chromosome loss

Thus far, we have shown that targeted chromosome loss can occur when using Cas9 to disrupt a desired gene, which predominantly occurs through end-joining DNA repair pathways. Tremendous effort has also been invested toward using Cas9 genome editing to correct a mutation or insert a gene by homology-directed repair (HDR). Since end-joining repair and HDR are divergent DNA repair pathways that involve different proteins, undergo different amounts of resection of the DSB ends, and occur in different stages of the cell cycle,^38^ we wanted to determine whether chromosome loss after Cas9-mediated genome editing also occurs during HDR. To explore this, we used Cas9 RNPs with a gRNA targeting *CD5* and various oligonucleotide HDR templates to integrate a short hemagglutinin (HA) tag in-frame with CD5.^39^ Successful generation of CD5+/HA+ cells via HDR peaked as high as ∼40% three or five after nucleofection (Fig. S5A). We performed ddPCR to quantify chromosome loss rates at both timepoints and observed similar levels of chromosome loss in *CD5*-targeted cells with an HDR template compared to *CD5*-targeted cells without an HDR template (Fig. S5B). Additionally, using CD5 and other T cell surface markers, we attempted to use fluorescence-activated cell sorting to enrich for cells without chromosome loss; however, we observed no reduction in chromosome loss (Supplementary Note 1).

### Pre-clinical CAR T cell generation results in chromosome loss

While CAR T cells and transgenic TCR T cells are currently manufactured using a retrovirus or lentivirus to semi-randomly integrate the retargeting transgene, researchers have also used Cas9 and HDR to precisely insert the transgene within the *TRAC* locus.^22, 39, 40^ This approach, which utilizes the native *TRAC* promoter to control expression of the CAR or retargeted TCR, offers advantages including uniform expression, minimal tonic signaling, and simultaneous disruption of the endogenous TCR. To explore whether chromosome loss occurs when generating pre-clinical CAR T cells via HDR, we nucleofected primary human T cells with Cas9 complexed with one of two *TRAC*-targeting gRNAs or a non-targeting gRNA. Just after nucleofection, recombinant adeno-associated virus 6 (AAV6) encoding a 1928(; CAR as an HDR template was added to the T cells (Fig. 5A and 5B).^41, 42^ Both the reduction of TCR expression and the gain of CAR expression were observed in two independent nucleofections; cells from one nucleofection were subjected to scRNA-seq at day four post-manufacturing, while cells from the other were subjected to scRNA-seq at day seven post-manufacturing. Overall rates of CAR integration via HDR were ∼34-69% (Fig. S5C). In all conditions that received a *TRAC*-targeting gRNA, regardless of day or whether an HDR template was present, we observed an enrichment in chromosome 14 loss compared to conditions with a non-targeting gRNA (Fig. 5C, S5D, and S5E). When chromosome 14 loss enrichment was normalized to editing efficacy, since day four and day seven CAR T cells were engineered independently, we observed a slight decrease in chromosome 14 loss enrichment over time (Fig. 5D). Together, these data suggest that chromosome loss is a general phenomenon that occurs in Cas9-edited T cells, regardless of the DNA repair pathway involved. Our findings also indicate that cells with chromosome loss are present among pre-clinical, Cas9-edited CAR T cells, highlighting the importance of understanding and mitigating this genotoxicity in the context of engineered T cell therapies.

**Figure 5:**
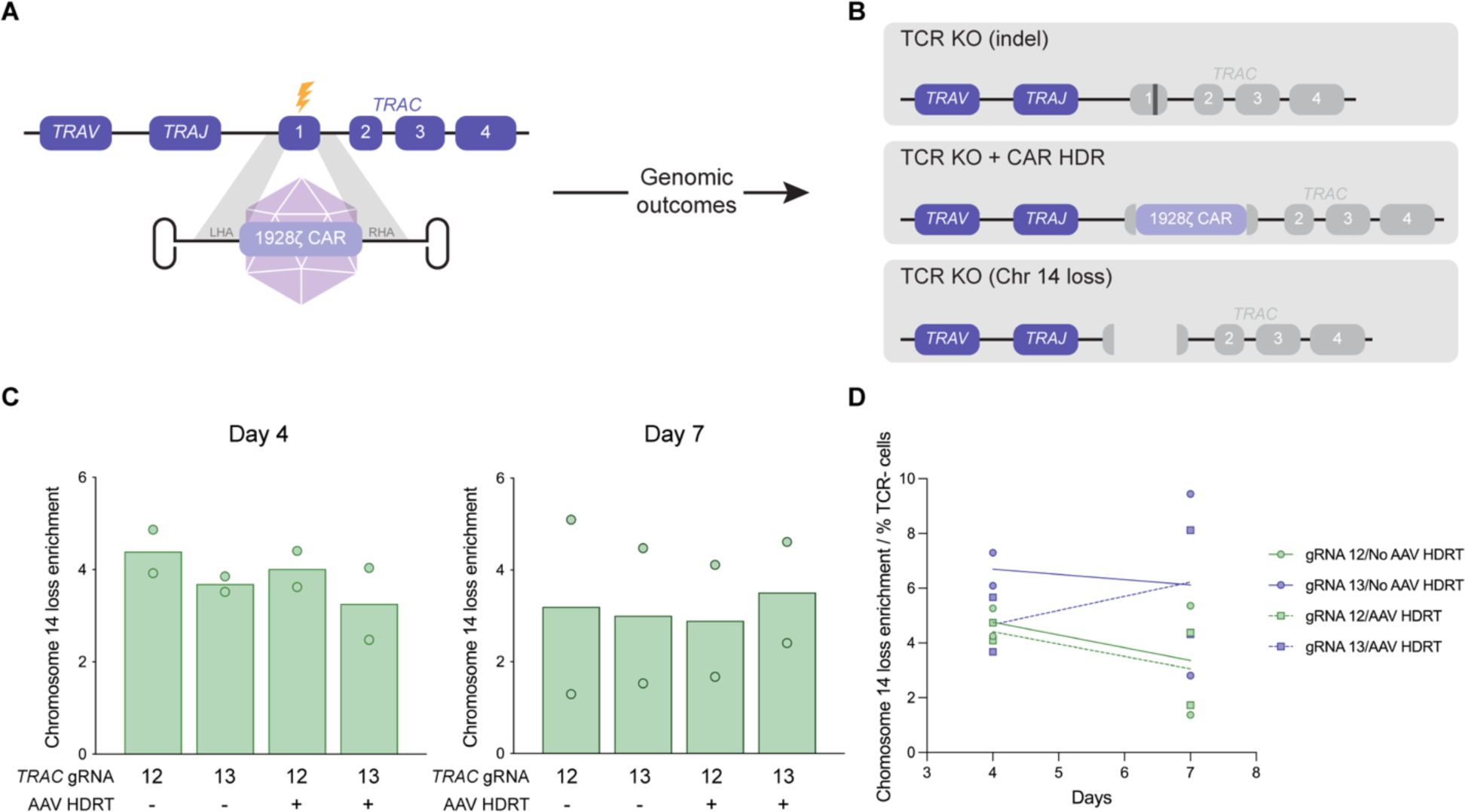
Pre-clinical CAR T cell production via homology-directed repair results in chromosome loss. **(A)** Strategy to generate CAR T cells via HDR with Cas9. AAV6 encoding a 1928(; CAR between left and right homology arms (LHA and RHA, respectively) serves as a template for HDR after Cas9 cleavage (yellow lightning bolt) of *TRAC*. **(B)** Three potential genomic outcomes after Cas9 HDR: indels that disrupt TCR expression (top), insertion of the CAR transgene that simultaneously disrupts TCR expression (middle), and chromosome loss that disrupts TCR expression (bottom). **(C)** Quantification of chromosome 14 loss enrichment across two *TRAC*-targeting gRNAs with or without an AAV HDR template from scRNA-seq (n = 2 biological donors). Two separate batches of CAR T cells were manufactured, before being subjected to scRNA-seq four or seven days after generation. Chromosome 14 loss enrichment was calculated relative to T cells treated with Cas9 and a non-targeting gRNA. **(D)** Chromosome 14 loss enrichment over time, normalized to Cas9 editing efficacy (n = 2 biological donors). Editing efficacy was determined by the percentage of TCR negative cells as measured via flow cytometry (see Extended Data Fig. 13c).

### Investigation of chromosome loss in Cas9-edited T cells from clinical trial patients

Our studies thus far have focused on *ex vivo* culturing of T cells; it is not yet known how these findings translate *in vivo*. We conducted a first-in-human phase 1 clinical trial (clinicaltrials.gov, trial NCT03399448) where Cas9 genome edited T cells were administered to patients with advanced, refractory cancer.^1^ Autologous T cells from three cancer patients were collected and electroporated with three different Cas9 RNPs, simultaneously targeting *TRAC*, *TRBC*, and *PDCD1*. These edited T cells were then transduced with a lentivirus encoding an HLA-A2*0201-restricted TCR specific to a peptide from the NY-ESO1 and LAGE-1 cancer antigens, resulting in engineered T cells (NYCE). NYCE cells were infused back into the patients and were found to be well-tolerated.

To investigate whether clinical manufacturing of a Cas9-edited adoptive T cell therapy results in levels of chromosome loss similar to those observed in our laboratory studies, we analyzed scRNA-seq data from NYCE cells of two patients at various timepoints throughout the clinical trial. Cells were collected from patient UPN35 prior to infusion and at day 28 post-infusion, while cells from patient UPN39 were collected prior to infusion as well as at days 10 and 113 post-infusion (Fig. 6A). Similar to our laboratory experiments, we inferred gene dosage at each of the target chromosomes (*TRAC*, Chr14; *TRBC*, Chr7; *PDCD1*, Chr2) and looked for partial and whole chromosome loss. Surprisingly, we observed extremely low levels of chromosome loss at the targeted chromosomes, which were not enriched compared to background levels at non-targeted chromosomes (Fig. 6B).

**Figure 6:**
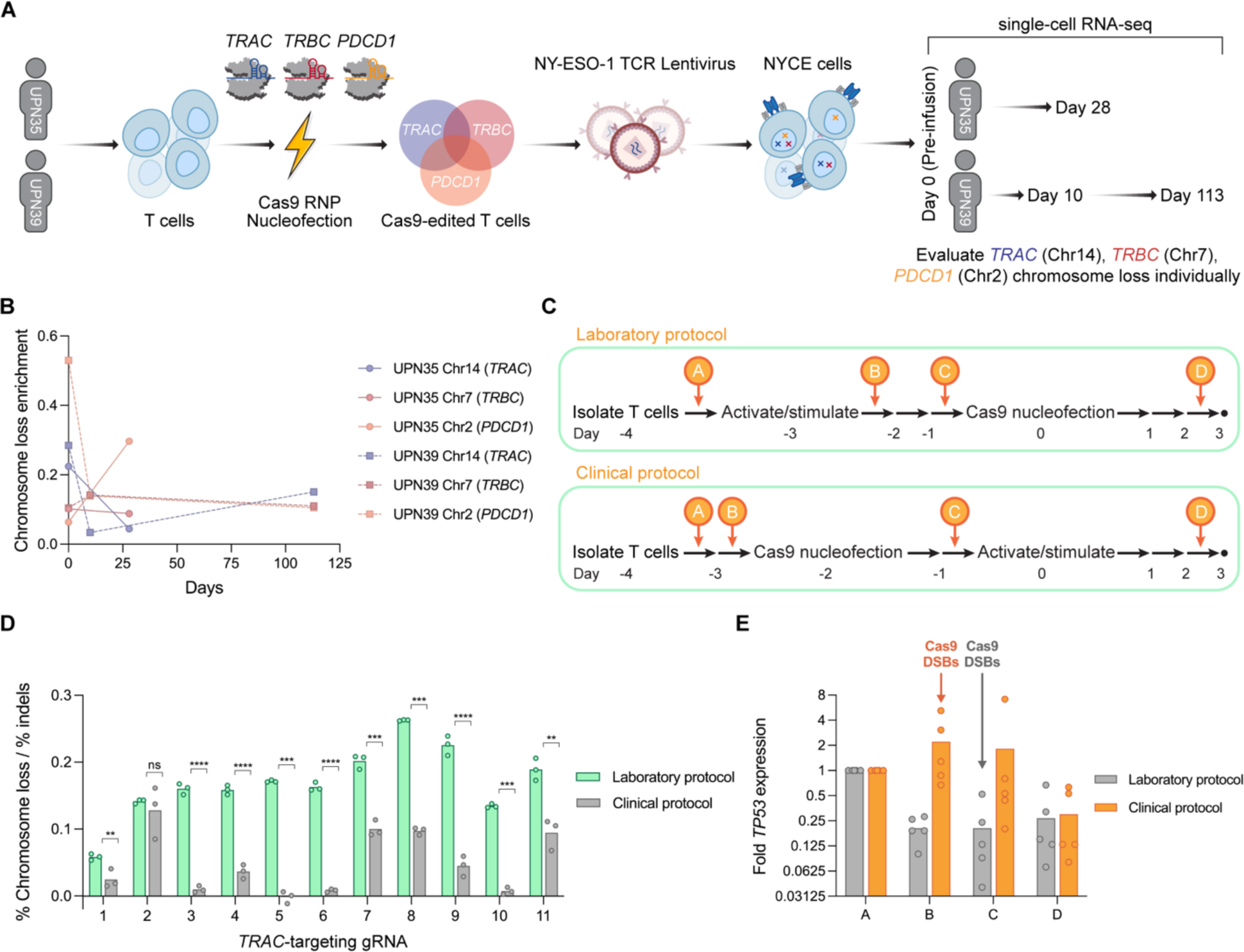
Clinical CRISPR-Cas9 genome editing protocol in patient T cells mitigates chromosome loss. **(A)** Strategy to investigate chromosome loss in two clinical trial patients with CRISPR-edited T cells. Two patients with refractory cancer had T cells isolated, nucleofected with *TRAC, TRBC,* and *PDCD1*-targeting Cas9 RNPs, and transduced with a lentivirus encoding an NY-ESO-1 TCR (NYCE cells). Cells were subjected to scRNA-seq prior to infusion (Day 0) and as well as at different time points post-infusion (Days 10, 28, and/or 113). **(B)** Chromosome loss enrichment on chromosome 14 (*TRAC*), chromosome 7 (*TRBC*), or chromosome 2 (*PDCD1*) at different timepoints for both patients. Enrichment was calculated relative to non-targeted chromosomes (all chromosomes but 2, 7, and 14). Day 0 represents NYCE cells prior to infusion, while other later timepoints represent NYCE cells that were collected after circulation *in vivo*. **(C)** Diagram of two different protocols for Cas9 genome editing of primary human T cells. The laboratory protocol (top) consisted of activating/stimulating cells prior to Cas9 nucleofection, and was used throughout this study. The clinical protocol (bottom) consisted of nucleofecting cells with Cas9 prior to activating/stimulating and is representative of our clinical trial. **(D)** Relative chromosome loss with 11 different *TRAC*-targeting gRNAs using the laboratory or clinical protocol in primary human T cells. Chromosome loss was normalized to the indel efficacy (see Extended Data Fig. 14b). *P*-values are from Welch’s unpaired t-tests and from left to right are 0.008320, 0.111695, 0.000052, 0.000076, 0.000159, 0.000073, 0.000125, 0.000087, 0.000073, 0.000050, and 0.001416. **(E)** Fold *TP53* mRNA expression during the laboratory or clinical protocols for Cas9 genome editing of primary human T cells (n = 5 biological donors). Data points are the mean of two technical replicates. X-axis letters correspond to timepoints in Fig. 6c. “Cas9 DSBs” represents the timepoints in the laboratory or clinical protocols where Cas9 was nucleofected into T cells to generate DSBs.

### Order of operations during Cas9 genome editing impacts chromosome loss

We wondered whether the discrepancy between the high rates of chromosome loss in our laboratory studies versus the low rates in our clinical trial could be attributed to the engineered T cell manufacturing protocol. In our laboratory studies, we activated and stimulated T cells prior to introducing Cas9 RNP and generating DSBs, while in our clinical trial we introduced Cas9 RNP and generated DSBs prior to activating and stimulating the T cells (Fig. 6C). We therefore performed Cas9 genome editing of *TRAC* in primary human T cells using our laboratory protocol (activation/stimulation followed by DSBs) and simulating our clinical protocol (DSBs followed by activation/stimulation) in parallel. Across all 11 *TRAC*-targeting gRNAs, we observed markedly lower levels of chromosome loss using our clinical protocol compared to our laboratory protocol (Fig. S6A). However, genome editing with the clinical protocol on average resulted in ∼14% lower indels compared to the laboratory protocol (Fig. S6B). To control for this difference, we normalized the rate of chromosome loss to the rate of indels generated per gRNA and still observed a statistically significant reduction in chromosome loss with our clinical protocol as compared to our laboratory protocol with 10 out of 11 gRNAs (Fig. 6D).

Previous studies have shown that p53, a key protein in cell cycle regulation and apoptosis, also regulates T cell activation. Downregulation of p53 is critical for murine T cell proliferation.^43^ Additionally, p53-mediated apoptosis has been shown as a mechanism for selecting against aneuploid cells.^44^ Therefore, we tested whether differences in manufacturing protocol influenced p53 levels, and how that related to the chromosome loss we observed. Expression of *TP53*, which encodes p53, was measured via RT-qPCR at different timepoints throughout both T cell genome editing protocols. Similar to what was observed in murine T cells, expression of p53 was lowered in both protocols after human T cell activation/stimulation (Fig. 6E). The mean *TP53* expression across five biological donors was >10 times higher immediately prior to Cas9-induced DSBs in our clinical protocol compared to our laboratory protocol (Fig. 6E). Thus, *TP53* expression during Cas9-induced DSBs inversely correlated with rates of chromosome loss in T cells between our two protocols. For our clinical protocol, the higher expression of this key DNA damage response factor during Cas9-induced DSBs could select against cells with chromosome loss and explain the dramatically lower rates we observed. Implementation of this modified protocol for Cas9 genome editing in T cells represents a simple adjustment that could substantially mitigate chromosome loss in future research and clinical studies.

## Discussion

In this study, we comprehensively investigated the frequency and consequences of Cas9-induced chromosome loss in primary human T cells, taking a genome-scale approach to understand what influences this phenomenon and investigating both pre-clinical and clinical T cell therapies. Targeting Cas9 across the *TRAC* locus, we estimated chromosome loss in ∼5-20% of cells depending on the gRNA. We discovered that Cas9-induced chromosome loss was a generalizable phenomenon; chromosome loss was observed across the genome in an average of 3.25% of T cells that were targeted by Cas9. These T cells showed detectable levels of chromosome loss over 2-3 weeks of *ex vivo* culture, though they displayed a fitness and proliferative disadvantage. These disadvantages could cause cells without chromosome loss to outgrow those with chromosome loss, explaining the gradual reduction in this chromosomal aberration measured over the multi-week culture. Importantly, we still detected chromosome loss in nearly all conditions at our final timepoint, and this 2-3 week timeframe is similar to current clinical adoptive T cell therapy manufacturing protocols.^1, 7^ This suggests that T cells with Cas9-induced chromosome loss could persist throughout *ex vivo* manufacturing and end up in the final product administered to patients. In addition, continued efforts aim to further shorten the engineered T cell manufacturing process, which has been shown to improve T cell activity and persistence but could result in higher levels of chromosome loss.^45^

To date, no study has investigated Cas9-induced chromosome loss in a clinical setting. In order to determine clinical significance, we generated CAR T cells using Cas9-mediated HDR, an approach being used in a growing number of clinical trials,^8, 9^ and found a significant enrichment in chromosome loss compared to non-targeted cells. We also investigated Cas9-edited T cells of two patients enrolled in a first-in-human phase 1 clinical trial. We previously reported detectable levels of Cas9-induced translocations, another unintended genomic aberration, in these patient T cells, though levels reduced to the limit-of-detection after *in vivo* circulation.^1^ Surprisingly, when we investigated patients’ T cells for chromosome loss, we saw no enrichment above background levels, marking two of the few cases where we did not find Cas9-induced chromosome loss.

Comparing the results from our laboratory experiments (where substantial chromosome loss was detected) and the clinical trial (where we did not observe chromosome loss above background levels), there were multiple technical differences in the parameters used for chromosome loss estimation (Supplementary Note 2). We tried to account for these differences by downsampling the CROP-seq screen dataset so that its parameters were similar to the clinical trial dataset, which was sparser in data (Fig. S6C). Even upon downsampling, our estimations of chromosome loss in the CROP-seq screen were comparable to the original complete dataset (Fig. S6D). This supports the conclusion that biological rather than technical reasons explain the dramatic difference in chromosome loss estimation.

We considered and eliminated multiple factors that might correlate with or potentially explain Cas9-induced chromosome loss including Cas9 binding orientation, gRNA sequence, chromatin accessibility, and targeted gene or chromosome. Instead, we found that introducing Cas9-mediated DSBs prior to T cell activation/stimulation, a protocol used in our clinical trial but not in our laboratory experiments, influenced this phenomenon by significantly diminishing chromosome loss. It is possible that high levels of transcription in activated T cells during our laboratory protocol may predispose cells to chromosome loss due to genome instability caused by active transcription.^46^ This effect could also be explained by levels of the DNA damage response protein p53 at the time of DSB generation, since we found *TP53* expression and chromosome loss were inversely correlated. Consistent with this hypothesis, a report in immortalized fibroblasts showed knockout of p53 increased Cas9-induced chromosomal truncations.^47^ For engineering T cells, using the manufacturing protocol in which cells are activated after delivery of Cas9 could become standard practice to minimize chromosome loss in the manufactured product. This protocol adjustment does not require novel equipment, modification of Cas9 or its gRNA, or additional cost, meaning it can be easily and immediately integrated into clinical practice.

Recently, several other Cas9-mediated chromosomal abnormalities such as translocations,^1^ large deletions,^48, 49^ loss of heterozygosity,^50^ and chromothripsis^51^ have been reported. Of these, only methods and technologies for limiting translocations have been demonstrated, including serial rather than simultaneous multiplexed genome editing,^52^ use of Cas12 nucleases,^53^ fusion of Cas9 to an exonuclease to limit repeat cleavage,^54^ or utilizing base editors that do not generate DSBs.^55^ Along with our modified clinical protocol, additional technologies could be developed to similarly mitigate chromosome loss.

CRISPR-based technologies that do not generate a DSB, such as base editors or epigenome editors, would likely avoid high levels of chromosome loss.^56, 57^ However, base editing can only modify one or a few nucleotides and epigenetic editing lacks permanence; neither of these technologies are ideal for permanent gene disruption or gene insertion. The use of CRISPR-Cas9 genome editing that creates DSBs is still highly advantageous and will continue to expand in clinical use. Therefore, mitigating genomic aberrations from DSBs, such as chromosome loss, will have substantial value to avoid potential genotoxicity in patients. Our comprehensive study suggests that although chromosome loss is a universal consequence of site-specific Cas9 genome editing, protocol adjustments and further exploration of underlying mechanisms can minimize its occurrence and impact.

### Limitations of the Study

To determine the generalizability of Cas9-induced chromosome loss, we performed a CROP-seq CRISPR screen targeting several genes on each somatic chromosome. The emergence of higher throughput scRNA-seq may allow this study to be expanded to a genome-wide screen in the future. Additionally, where possible we selected highly active and specific gRNAs from previously reported studies to include in our CROP-seq library. However, since we cannot reliably measure genome editing efficacy in our pooled format, it is possible that gRNAs with low chromosome loss detected simply had low cleavage activity. Finally, sequencing quality and computational gene calling varied between experiments (Supplementary Note 2). Since this could influence our chromosome loss measurements, we primarily displayed relative chromosome loss enrichment, which was normalized to a non-targeting gRNA or untreated sample, rather than absolute chromosome loss.

### Star Methods

#### Cell culture

Primary adult peripheral blood mononuclear cells (PBMCs) were obtained as cryopreserved vials from Allcells Inc. CD3+ T cells were isolated from PBMCs using EasySep Human T Cell Isolation Kits (StemCell Technologies) according to the manufacturer’s instructions. Isolated CD3+ T cells were cultured in X-Vivo 15 medium (Lonza) with 5% fetal bovine serum (FBS) (VWR), 50 μM 2-mercaptoethanol, and 10 mM N-acetyl L-cysteine (Sigma-Aldrich). One day post-isolation, CD3+ T cells were activated and stimulated with a 1:1 ratio of anti-human CD3/CD28 magnetic Dynabeads (Thermo Fisher) to cells, as well as 5 ng/mL IL-7 (PeproTech), 5 ng/mL IL-15 (R&D Systems), and 300 U/mL IL-2 (PeproTech) for three days. After the initial activation and stimulation, magnetic beads were removed and T cells were cultured in medium with 300 U/mL IL-2. Medium was replaced every other day and T cells were maintained at a density of ∼0.5-1×10^6^ cells/mL.

For CAR T cell experiments, primary adult PBMCs were obtained as Leukopaks (StemCell Technologies) from deidentified healthy donors and cryopreserved in RPMI medium supplemented with 20% human serum and 10% DMSO. T cells were isolated as described before and cultured in X-Vivo 15 medium supplemented with 5% human serum, 5 ng/mL IL-7 (Miltenyi Biotec), and 5 ng/mL IL-15 (Miltenyi Biotec). Immediately after isolation, T cells were stimulated for two days with anti-human CD3/CD28 magnetic Dynabeads (Thermo Fisher) using a 1:1 bead-to-cell ratio.

#### Cas9 ribonucleoprotein nucleofection

100 pmol Alt-R crRNA and 100 pmol Alt-R tracrRNA (IDT) were diluted in IDT Duplex Buffer, incubated at 90° C for 5 min, and then slow cooled to room temperature (Supplementary Tables 1, 7-8). 50 pmol *S. pyogenes* Cas9 V3 (IDT) was diluted in RNP buffer (20 mM HEPES, 150 mM NaCl, 10% Glycerol, 1 mM MgCl_2_, pH 7.5). Cas9 and duplexed gRNA (1:2 molar ratio) were incubated at 37° C for 15 min. Primary human T cells were washed once with PBS (-/-) before 250,000 cells were resuspended in P3 Buffer (Lonza). 50 pmol Cas9 RNP was added to the cells before nucleofection in a Lonza 4D-Nucleofector with pulse code EH-115. X-Vivo 15 medium with 300 U/mL IL-2 was added to the nucleofected cells before a 30 min recovery at 37° C. Nucleofected T cells were plated at a density of ∼0.5-1×10^6^ cells/mL in 96-well U-bottom plates.

#### TCR flow cytometry

T cells were resuspended in Cell Staining Buffer (BioLegend) with Ghost Dye Red 780 (1:1,000, TonboBio) and anti-human TCR α/ý Brilliant Violet 421 (1:100, BioLegend). Cells were stained for 30 min at 4° C in the dark. After staining, cells were washed with Cell Staining Buffer and analyzed on a Thermo Fisher Attune NXT Flow Cytometer with an autosampler. Over 20,000-100,000 cells were routinely collected and analyzed with FlowJo.

#### Next-generation sequencing of TRAC genome editing

Genomic DNA from T cells was extracted with QuickExtract DNA Extraction Solution (Lucigen) by incubating resuspended cells for 10 minutes at room temperature before heating lysates at 65° C for 20 minutes then 95° C for 20 min. The region of *TRAC* containing the Cas9 target site was amplified from genomic DNA with Q5 High-Fidelity DNA polymerase (NEB) to add universal adaptors (Supplementary Table 2). Amplicons were cleaned with SPRIselect beads (Beckman Coulter) before a second round of PCR was performed to add unique i5 and i7 Illumina indices to each sample. Subsequent amplicons were cleaned again, and libraries were sequenced on an Illumina iSeq 100 (2×150 bp). FASTQ files were trimmed, merged, and analyzed for indels with CRISPResso2 (crispresso2.pinellolab.org).^58^ For non-targeting conditions, sequencing from cells receiving Cas9 and a non-targeting gRNA was used as input to check for indels at a given region.

#### Single-cell RNA sequencing with MULTI-seq barcoding

Four days post-nucleofection, T cells were labeled via MULTI-seq as previously described.^15^ Briefly, a lipid-modified oligonucleotide (LMO) was combined with a unique oligonucleotide barcode (Supplementary Tables 3 and 6) at a 1:1 molar ratio in PBS (-/-). 500,000 cells or fewer were washed twice with PBS (-/-) and then resuspended in PBS (-/-). The LMO/barcode solution was mixed with each cell suspension and incubated on ice for 5 minutes before addition of a co-anchor LMO and incubation on ice for an additional 5 minutes. Cold 1% BSA in PBS (-/-) was added to sequester free LMOs before washing cells twice with cold 1% BSA in PBS (-/-). Uniquely labeled cells were pooled in equal numbers in 1% BSA in PBS (-/-) to a final concentration of ∼1,600 cells/mL. 10x Genomics Chromium Next GEM Single Cell 3’ Gene Expression Kits (v3.1) were utilized according to the manufacturer’s instructions, with the following modifications. Lanes of a standard Chromium chip were “super loaded” with ∼50,000 cells to yield a target cell recovery of ∼25,000 cells. During the cDNA amplification, 1 μL of a 2.5 μM MULTI-seq primer (see McGinnis et al.) was added. Supernatant from the cDNA bead cleanup was saved because it contained the MULTI-seq barcode amplicon. Supernatants were further cleaned by addition of SPRIselect beads and isopropanol with a conventional magnetic bead cleanup protocol. 3.5 ng of each cleaned amplicon was used in a PCR reaction to add sequencing indices; the reactions included KAPA HiFi HotStart ReadyMix (Roche), a unique i5 primer, and a unique RPI i7 primer (Supplementary Table 3). The PCR reactions were cleaned with SPRIselect beads before final library QC. Gene expression and MULTI-seq barcode libraries were pooled 6:1 (molar ratio) and sequenced on an Illumina NovaSeq 600 S1 Flow Cell.

#### Single-cell RNA sequencing analysis

Cell Ranger (v7.0) was used to process Chromium single cell data. cellranger count was performed with the parameters--r1-length=28 and--r2-length=90. For the first *TRAC*-targeting experiment the--force-cells parameter was set to 15,000 cells. To demultiplex the different pools of cells using the MULTI-seq barcode, cellranger multi was performed. The results from the different pools were aggregated using cellranger aggr. The results from cellranger were parsed with scanpy and converted to a h5ad file format. The demultiplexed results were added as metadata to the h5ad file.

For the CROP-seq screen, we counted the number of reads within each cell aligning to each of the gRNAs used in the screen. To determine which cells were targeted by a single, unique gRNA, we tested whether the gRNA with the highest number of reads had significantly more reads than the second highest. Specifically, let *c*_1_and *c*_2_ be the number of reads for the first and second most common gRNAs in a cell, respectively. To test whether *c*_1_ is significantly greater than *c*_2_, we calculated a *P*-value based on a binomial distribution with parameters *n* = *c*_1_ + *c*_2_ and *p* = 0.5 (i.e. *x*∼*B*(*c*_1_ + *c*_"2_ 0.5)). If the probability of *x* ≥ *c*_1_ was smaller than 0.05, the cell was determined to be transduced by a single gRNA.

#### Quantification of chromosome loss from scRNA-seq

To assess the dosage of each gene in each cell, inferCNV of the Trinity CTAT project (https://github.com/broadinstitute/infercnv) was executed in R (version 4.1) with default parameters over the h5ad dataset created by cellranger (see previous section) for every scRNA-seq dataset.^16^ Each cell with a gRNA was labeled as a “treatment” and each cell with a non-targeted gRNA was labeled as a “control.” To successfully run inferCNV for the CROP-seq screen, inferCNV was performed in multiple batches of 30,000 cells.

The output of inferCNV was the estimated dosage for each gene; according to the software’s specifications, values below 0.95 were considered loss of at least one copy of the gene. inferCNV values for each gene were binarized as <0.95 or ≥ 0.95. Each chromosome in each cell was then searched for the interval between two genes that maximizes the difference between the average binarized inferCNV values on either side of the interval. This interval was the candidate breakage point for a particular chromosome in a cell.

We used the inferCNV values for all genes on a given chromosome within each cell (with respect to each of the 22 somatic chromosomes) to estimate the loss status of that chromosome. Specifically, we estimated whether there was 1) no chromosome loss, 2) whole chromosome loss, or 3) partial chromosome loss. If at least 70% of a minimum 150 genes to either the left or right of the candidate breakage point were below the 0.95 threshold, but less than 70% of the genes on the other side were below the threshold, the cell was labeled as partial chromosome loss for that chromosome. Otherwise, if at least 70% of all the genes throughout the entire chromosome were below the threshold, the cell was labeled as having whole chromosome loss for that chromosome. If neither were true, the cell was labeled as having no chromosome loss for that chromosome.

Downsampling of the CROP-seq screen dataset was performed to assess the dependency of chromosome loss enrichment on the number of genes in the inferCNV output. To do this, 4,000 or 1,000 genes contained within the 9,639 gene output of the full CROP-seq inferCNV output were randomly sampled. Our chromosome loss calling pipeline, as described above, was then performed on these downsampled datasets.

#### Droplet digital quantitative PCR

Genomic DNA was collected from T cells at different times post-nucleofection with QuickExtract DNA Extraction Solution, identical to as described earlier. The ddPCR setup was similar to what has been previously described.^37^ For multiplexed ddPCR, two ∼200 bp amplicons for each target gene were designed (Supplementary Tables 4, 7, and 8). Amplicon 1 was located proximal to the centromere and utilized a hexachlorofluorescein-labeled (HEX) oligonucleotide probe (PrimeTime qPCR probes, Zen double quencher, IDT). Amplicon 2 was located ∼100-200 bp away from amplicon 1, was distal relative to the centromere, and utilized a 6-fluorescein-labeled (FAM) oligonucleotide probe (PrimeTime qPCR probes, Zen double quencher, IDT). Amplicon 1 served as a control, which should be unaffected by Cas9 genome editing or chromosome loss and would signal whether the gene of interest was in a given droplet. Amplicon 2 spanned the Cas9 target site, with the probe located ∼30-60 bp away from the cleavage site. If the target site was not successfully repaired after Cas9 cleavage, amplicon 2 would not be able to be amplified and the FAM probe would be unable to dissociate from its quencher. ddPCR reactions were assembled with ddPCR Supermix for Probes (No dUTP, Bio-Rad), 900 nM of each primer, 250 nM of each probe, and 10-30 ng of genomic DNA. Droplets were formed using a Bio-Rad QX200 Droplet Generator following the manufacturer’s instructions before thermal cycling. The following day, ddPCR droplets were analyzed on a Bio-Rad QX200 Droplet Reader. Data were analyzed with the QX Manager Software (Bio-Rad), and thresholds were set manually based on wells with untreated samples. The percentage of alleles with chromosome loss was calculated based on droplets that had the target amplicon 1 (HEX+) but were unable to produce the neighboring amplicon 2 (FAM-). The equation utilized is as follows: [inlineequation1]

#### Genome-scale CROP-seq CRISPR screen design

The CROP-seq library was designed to contain multiple gRNAs that target multiple genes on every chromosome. When possible, validated gRNA sequences from previous publications were utilized (Supplementary Table 5). The gRNA library was ordered as an oPool oligo pool (IDT) and Golden Gate cloned into a custom CROP-seq vector that co-expressed GFP. To analyze the library, primers were used to amplify the gRNA spacer from either the plasmid library or genomic DNA library before sequencing on an Illumina iSeq. MAGeCK (https://sourceforge.net/p/mageck/wiki/Home/) was used to quantify the representation of each gRNA in the library.^59^

#### CROP-seq CRISPR screen lentiviral production

For lentivirus production, Human Embryonic Kidney 293T (HEK293T) cells were cultured in Dulbecco’s Modified Eagle Medium (Gibco) with 10% FBS and 1% penicillin/streptomycin (Gibco). HEK293Ts were transfected at 70-90% confluency with 10 μg CROP-seq gRNA plasmid, 10 μg Gag-pol expression plasmid (psPax2, gift from Didier Trono, Addgene plasmid #12260), and 1 μg pCMV-VSV-G plasmid (gift from Bob Weinberg, Addgene plasmid #121669) using polyethylenimine (PEI, Polysciences Inc.) at a 3:1 PEI:plasmid ratio. Approximately 6-8 hours after transfection, the medium was aspirated from cells and replaced with Opti-Mem (Gibco). Supernatant containing lentivirus was collected 48 hours after transfection, the medium was replaced, and medium was collected once more after an additional 48 hours. Viral supernatants were filtered through a 0.45 μm PES membrane bottle top filter (Thermo Fisher) and then concentrated with Lenti-X Concentrator (Takara) according to the manufacturer’s instructions. Purified and concentrated lentivirus was used immediately or stored at −80° C. Lentivirus was titered by counting the number of initially transduced cells, adding serial dilutions of lentivirus to primary human T cells, and measuring the percentage of GFP+ cells after three days (only in conditions with <30% GFP+ cells to ensure a majority were single transduction events).

#### CROP-seq CRISPR screen

For the CROP-seq screen, primary human T cells were isolated and stimulated as stated previously. 24 hours after stimulation, lentivirus was added to the cells at a multiplicity of infection (MOI) of ∼0.3. MOI was confirmed via flow cytometry two days later, at day three post-stimulation. Dynabeads were then removed from T cells and Cas9 was nucleofected as stated previously. For the full CROP-seq library experiment, four days post-nucleofection, T cells were subject to fluorescence-activated cell sorting (FACS) on a Sony SH800S cell sorter to enrich for GFP+ cells. Genomic DNA was harvested from a small number of cells, as previously described, to assess the library representation. The rest of the live/GFP+ cells were arbitrarily divided into six pools and subject to MULTI-seq barcoding and 10x Genomics scRNA-seq, as previously described. The CROP-seq gRNA was enriched from the resulting cDNA similar to what has been previously described (Supplementary Table 6).^20^ Briefly, 25 ng of cDNA was added to eight separate KAPA HiFi HotStart ReadyMix PCR reactions and amplified for 10 cycles with an annealing temperature of 65° C to enrich for the gRNA. Individual PCR reactions were pooled together and cleaned with SPRIselect beads. 8 μL of cleaned PCR1 product was added to a second KAPA HiFi HotStart ReadyMix PCR reaction and amplified for 10 cycles with an annealing temperature of 65° C to add Illumina sequencing adaptors. Gene expression, MULTI-seq barcode, and CROP-seq enrichment libraries were sequenced on an Illumina NovaSeq 600. Multiple iterations of library sequencing were concatenated to achieve the desired sequencing depth.

#### Strand and MMEJ analyses

Each gRNA was mapped using the GRCh38 genome assembly. A two-sided Fisher’s Exact Test was performed to determine whether gRNAs binding distal or proximal to the centromere, relative to the gRNA spacer sequence, affected chromosome loss. MMEJ analyses were performed using inDelphi (indelphi.giffordlab.mit.edu).^60^ The cell type was set to K562s and the MMEJ strength was measured for each unique gRNA sequence.

#### Differential gene expression analysis

To identify genes differentially expressed between cells with and without chromosome loss, we used the memento algorithm with default parameters (capture rate = 0.07).^61^ We tested each gene for differential expression with respect to each of the 22 somatic chromosomes separately, and only reported genes that were consistently over-or underexpressed across most chromosomes. We further ensured that the differences we observed were specific to the tested gene and were not the result of overall lower or greater levels of gene expression in cells with chromosome loss. To do this we accounted for the total expression of transcripts sharing the same chromosome as the tested gene by including the overall count of all transcripts on that chromosome as a covariate. This covariate was defined with respect to the chromosome containing the gene tested for differential expression and not with respect for the chromosome determining the two compared groups (cells with or without loss of that chromosome). Since memento supports only discrete covariates, we discretized the total transcript count into 10 decile bins.

We corrected the results of memento for multiple testing using FDR over the combined set of all tested genes over all tested chromosomes. We considered a gene to be statistically significant with respect to a chromosome (i.e. the gene to be over-or underexpressed in cells losing that chromosome) if its corrected *P*-value was below 0.05. Accordingly, we assigned the significance status of that gene-chromosome combination to be 1, 0, or −1 if it was significantly overexpressed, not significant, or under expressed, respectively. We then assigned each gene a total score between −22 and 22 by summing the significance status of that gene with respect to each of the 22 somatic chromosomes. 613 genes obtained a total score ≥5 and were considered the top overexpressed genes, while 590 obtained a total score ≤ −5 and were considered the top underexpressed genes.

We identified pathways enriched among the 613 top overexpressed genes by searching through the pathway terms defined in the KEGG database using the GSEApy Python package.^62, 63^

#### Cell cycle analysis

Cell cycle states were defined using data and methods as previously described.^64^

#### Epigenetic analyses

Datasets (ENCFF233TCT, ENCFF055FYI, and ENCFF129GAM) corresponding to activated T cells from a male donor (43 years old) were selected from the ENCODE Portal (www.enccodeproject.org).^65,66^ The presence of open chromatin from ATAC-seq data and the location of epigenetic marks (H3K9me3 and H3K36me3) from ChIP-seq data were determined within a 75 bp window around the GRCh38 coordinates of each gRNA and a *P*-value <10^-5^, according to best practices.^67^

#### T cell proliferation tracking

After isolation and stimulation, primary human T cells were nucleofected with Cas9 RNPs identical to what was described earlier. Immediately after nucleofection recovery, cells were pelleted and resuspended in 5 μM CellTrace Violet (Invitrogen). Cells were incubated in CellTrace Violet for 20 min at 37° C, prior to diluting in 4x volume of complete X-Vivo 15 medium to absorb unbound dye and incubating again for 5 min at 37° C. Cells were pelleted and resuspended in complete X-Vivo 15 with 300 U/mL IL-2. T cells were passaged every other day to refresh medium and maintain a density of ∼0.5-1×10^6^ cells/mL. Cells were sorted on a BD FACSAria II to obtain the approximate bottom and top quartile of cells according to CellTrace Violet signal.

#### Cas9-mediated CD5 homology-directed repair

HA tag insertion was achieved with either a single-stranded DNA HDR template (ssDNA HDRT), a double-stranded DNA HDR template (dsDNA HDRT), or a single-stranded DNA HDR template with Cas9 target sequences (ssCTS HDRT) (Supplementary Note 3).^39^ Equimolar HDRT oligonucleotides were diluted in IDT Duplex Buffer, heated to 95° C for 5 minutes, then allowed to slow cool to room temperature. 100 pmol of HDRT was added to Cas9 RNP nucleofections of primary human T cells, identical to as described above. Cells were analyzed on a Thermo Fisher Attune NXT Flow Cytometer with an autosampler, identical to as described above, except with the antibodies anti-human CD5 (UCHT2)-PE (1:100, Invitrogen) and anti-HA (6E2)-AF647 (1:100, Cell Signaling Technology). Over 20,000-100,000 cells were routinely collected and analyzed with FlowJo.

#### Fluorescence-activated cell sorting of CD5, CD81, and CD3E

Primary human T cells were nucleofected with Cas9 RNPs targeting *CD5* identical to what was described before. Seven days post-nucleofection, cells were stained with anti-human TCR α/ý Brilliant Violet 421 (BioLegend), anti-human CD5 (UCHT2)-PE (Invitrogen), anti-human CD81 (5A6)-FITC (BioLegend), and anti-human CD3E (SK7)-APC (Invitrogen), all at a 1:100 dilution. Cells were sorted on a BD FACSAria II to isolate different populations (Supplementary Note 1).

#### CAR adeno-associated virus production

An AAV transgene plasmid encoding the inverted terminal repeats, a 1928(; CAR, a truncated human EGFR (EGFRt) tag, and *TRAC* homology arms for HDR was used as previously described (Supplementary Note 4).^41^ The AAV plasmid was packaged into AAV6 by transfection of HEK293T cells together with pHelper and pAAV Rep-Cap plasmids using PEI. The AAVs were purified using iodixanol gradient ultracentrifugation. The titration of the AAV was determined by quantitative PCR on DNaseI (NEB) treated and proteinase K (Qiagen) digested AAV samples, using primers against the left homology arm. The quantitative PCR was performed with SsoFast EvaGreen Supermix (Bio-Rad) on a StepOnePlus Real-Time PCR System (Applied Biosystems).

#### CAR T cell production

gRNAs targeting exon 1 of the *TRAC* locus (*TRAC* gRNA 12), the intron preceding the *TRAC* locus (*TRAC* gRNA 13), or a non-targeting control gRNA were purchased from Synthego and resuspended in TE buffer (Supplementary Table 9). Cas9 RNP was generated by incubating 60 pmol of Cas9 protein with 120 pmol sgRNA. T cells were counted, resuspended in P3 buffer at 2×10^6^ per 20 μL, mixed with 3 μL of RNPs and added to a 96-well nucleofection plate. Cells were electroporated using a Lonza 4D-Nucleofector 96-well unit with the EH-115 protocol and immediately recovered by adding pre-warmed X-Vivo 15 medium without human serum. Recombinant AAV6 encoding the HDR template was added to the culture 30 to 60 min after nucleofection at an MOI of 10^5^, and incubated with the cells overnight. The day after the nucleofection and transduction, edited cells were resuspended in medium and expanded using standard culture conditions, keeping a density of 10^6^ cells/mL. TCR disruption and CAR HDR efficiency was evaluated by flow cytometry by staining the TCR with anti-TCRα/ý (BW242/412)-PE (1:50, Miltenyi) and the CAR with goat anti-mouse IgG (H+L) AlexaFluor 647 Fab (1:100, Jackson ImmunoResearch).

#### CAR T cell scRNA-seq

CAR T cells were harvested at two time points after independent nucleofections (day four and day seven post-nucleofection). TotalSeq-A0251-1 anti-human Hashtag reagents (BioLegend) were used to label different cell conditions. For the experiment, 500,000 cells from each condition were labeled with the hash antibodies in Cell Staining Buffer at 4° C for 30 min. After labeling, cells were washed three times with Cell Staining Buffer at 4° C and then resuspended in PBS (-/-) containing 0.04% BSA. Labeled cells were pooled and 50,000 cells were “super loaded” into four lanes (two lanes for day four samples and the other two lanes for day seven samples) of a 10X Chromium Single-Cell G Chip. A 10x Genomics Chromium Next GEM Single Cell 3’ Gene Expression Kit (v3.1) was utilized according to the manufacturer’s instructions, and the subsequent library was sequenced on an Illumina NovaSeq 600 S4 Flow Cell.

#### Laboratory versus clinical T cell manufacturing

For the laboratory protocol, T cells were activated and stimulated identical to what was described earlier. After nucleofection of Cas9 RNP, T cells were cultured in X-Vivo 15 medium with 5 ng/mL IL-7, 5 ng/mL IL-15, and 300 U/mL IL-2.

For the clinical protocol, we followed a protocol similar to what was used in our phase 1 clinical trial^1^. After T cell isolation, cells were cultured in X-Vivo 15 medium with 5 ng/mL IL-7 and 5 ng/mL IL-15 for two days. Non-activated T cells were nucleofected with 50 pmol Cas9 RNP using a Lonza 4D-Nucleofector with pulse code EH-115. After nucleofection, cells were incubated in X-Vivo 15 medium with 5 ng/mL IL-7 and 5 ng/mL IL-15 for a 30 min recovery at 37° C. Nucleofected T cells were plated at a density of ∼0.5-1×10^6^ cells/mL in 96-well U-bottom plates. Two days after nucleofection, cells were counted and activated/stimulated with a 1:1 ratio of anti-human CD3/CD28 magnetic Dynabeads to cells, as well as 5 ng/mL IL-7, 5 ng/mL IL-15, and 300 U/mL IL-2 for an additional three days.

#### T cell RT-qPCR

During the laboratory or clinical T cell manufacturing protocols, 500,000 cells were periodically pelleted and resuspended in TRIzol (Invitrogen). RNA was isolated via phenol-chloroform extraction, precipitated by addition of isopropanol, washed with 75% ethanol, and resuspended in nuclease-free water. Isolated RNA was treated with TURBO DNase (Invitrogen) and SUPERase-In RNase inhibitor (Thermo Fisher) for 30 min at 37° C before addition of DNase Inactivation Reagent according to the manufacturer’s instructions. DNA-free RNA underwent cDNA synthesis using SuperScript III Reverse Transcriptase (Invitrogen) and Random Primers (Promega) according to the manufacturer’s instructions. qPCRs were performed with the resulting cDNA using iTaq Universal SYBR Green Supermix (Bio-Rad) on a Bio-Rad CFX96 Real-Time PCR Detection System (Supplementary Table 10). *TP53* expression levels were normalized to the expression levels of the housekeeping gene *GAPDH,* and to timepoint A (where the laboratory and clinical protocols start identically) using the ýýCt method.

## Supplemental Information

## Acknowledgements

The authors would like to thank Jennifer Hamilton, Matthew Kan, Elizabeth Abby Stahl, and David Colognori for thoughtful discussion throughout this project. We thank Christopher McGinnis and Zev Gartner for providing the MULTI-seq reagents; Yoon Gi Justin Choi (QB3 Functional Genomics Laboratory) for assistance with the scRNA-seq; Scarleth Chalen Ulloa for assistance with the FACSAria; Netra Krishnappa (IGI Center for Translational Genomics), Carrianne Miller (QB3 Genomics Sequencing Laboratory), and Eric Chow (UCSF Center for Advanced Technology) for assistance with NGS; MinCheol Kim for assistance with the differential gene expression analysis; Francois Aguet for assistance with the cell cycle analysis; and Brian Shy for assistance with the CD5 HDR assay. We thank the patients, their families, and the clinical study team at the Abramson Cancer Center Clinical Research Unit for their involvement in the clinical trial. We thank all members of the Satpathy, Cate, Eyquem, Fraietta, June, Chang, Ye, and Doudna laboratories for their input and advice. C.A.T. is supported by a National Institutes of Health (NIH) Ruth L. Kirschstein National Research Service Award F31 Pre-Doctoral Fellowship (National Heart, Lung, and Blood Institute, F31HL156468-01) and the Siebel Scholarship (Siebel Foundation). B.Y. is supported by the Parker Institute for Cancer Immunotherapy Parker Bridge Fellowship and the V Foundation. This project was supported by the Parker Institute for Cancer Immunotherapy (A.T.S., J.E., C.H.J., H.Y.C., and C.J.Y.), the NIH National Institute of General Medical Sciences (R01-GM065050, J.H.D.C.), the NIH National Cancer Institute (R35CA209919, H.Y.C), the NIH Centers for Excellence in Genomic Science (RM1HG007735, H.Y.C; RM1HG009490, J.A.D.), the Howard Hughes Medical Institute (H.Y.C. and J.A.D.), the NIH National Institute of Arthritis and Musculoskeletal and Skin Diseases (R01AR071522, C.J.Y.), the NIH National Institute of Allergy and Infectious Diseases (R01AI136972, C.J.Y.), the Chan Zuckerberg Initiative and Chan Zuckerberg Biohub (C.J.Y.), and the NIH Somatic Cell Genome Editing Program of the Common Fund (U01AI142817-02, J.A.D.).

## Author Contributions

C.A.T., J.H.D.C., and J.A.D. conceived the study with subsequent input from N.B., R.B., and C.J.Y. C.A.T., B.H., C.C., J.L., and Y.S. performed the T cell experiments and generated the laboratory scRNA-seq libraries. C.R.H. provided technical support on the clinical protocol. N.B., R.B., M.T., and T.M. analyzed the laboratory scRNA-seq data. K.R.P. and Y.Q. generated the clinical trial scRNA-seq libraries. M.T., T.M., and B.Y analyzed the clinical trial scRNA-seq data. A.T.S., E.A.S., J.H.D.C., J.E., J.A.F., C.H.J., H.Y.C., C.J.Y., and J.A.D. supervised the project. C.A.T., N.B., C.J.Y., and J.A.D. wrote the manuscript with input from all other authors.

## Declaration of Interests

C.A.T, J.A.D., and the Regents of the University of California have patents pending or issued related to the use of CRISPR genome editing technologies. R.B. is an employee of BioMarin Pharmaceutical Inc., J.L. is an employee of Altos Labs, and K.R.P. is a co-founder and employee of Cartography Biosciences. A.T.S is a co-founder of Immunai and Cartography Biosciences. A.T.S. has received research support from Arsenal Biosciences, Allogene Therapeutics, and 10x Genomics. J.H.D.C. is a co-founder of Initial Therapeutics. J.E. is a co-founder of Mnemo Therapeutics, a scientific advisory board member of Cytovia Therapeutics, and a consultant for Casdin Capital, Resolution Therapeutics, IndeeLabs, and Treefrog Therapeutics. J.E. has received research support from Cytovia Therapeutics, Mnemo Therapeutics, and Takeda Pharmaceutical Company. J.A.F has received research support from Tmunity. C.H.J. and the University of Pennsylvania have patents pending or issued related to the use of gene modification in T cells for adoptive T cell therapy. C.H.J. is a co-founder of Tmunity. H.Y.C. is a co-founder of Accent Therapeutics, Boundless Bio, Cartography Biosciences, and Orbital Therapeutics, and an advisor to 10x Genomics, Arsenal Biosciences, Chroma Medicine, Spring Discovery, and Vida Ventures. C.J.Y is a co-founder of Survey Genomics, and a scientific advisory board member of Related Sciences and Immunai. C.J.Y. is a consultant for Maze Therapeutics, TReX Bio, ImYoo, and Santa Ana Bio. C.J.Y. has received research support from the Chan Zuckerberg Initiative, Chan Zuckerberg Biohub, Genentech, BioLegend, ScaleBio, and Illumina. J.A.D. is a co-founder of Editas Medicine, Intellia Therapeutics, Caribou Biosciences, Mammoth Biosciences, and Scribe Therapeutics, and a scientific advisory board member of Intellia Therapeutics, Caribou Biosciences, Mammoth Biosciences, Scribe Therapeutics, Vertex Pharmaceuticals, eFFECTOR Therapeutics, Felix Biosciences, The Column Group, Inari, and Isomorphic Labs. J.A.D. is the Chief Science Advisor at Sixth Street, an advisor at Tempus, and a Director at Johnson & Johnson and Altos Labs. J.A.D. has sponsored research projects through Biogen, Pfizer, Apple Tree Partners, Genentech, and Roche. All other authors declare no competing interests.

---

**Figure S1:**
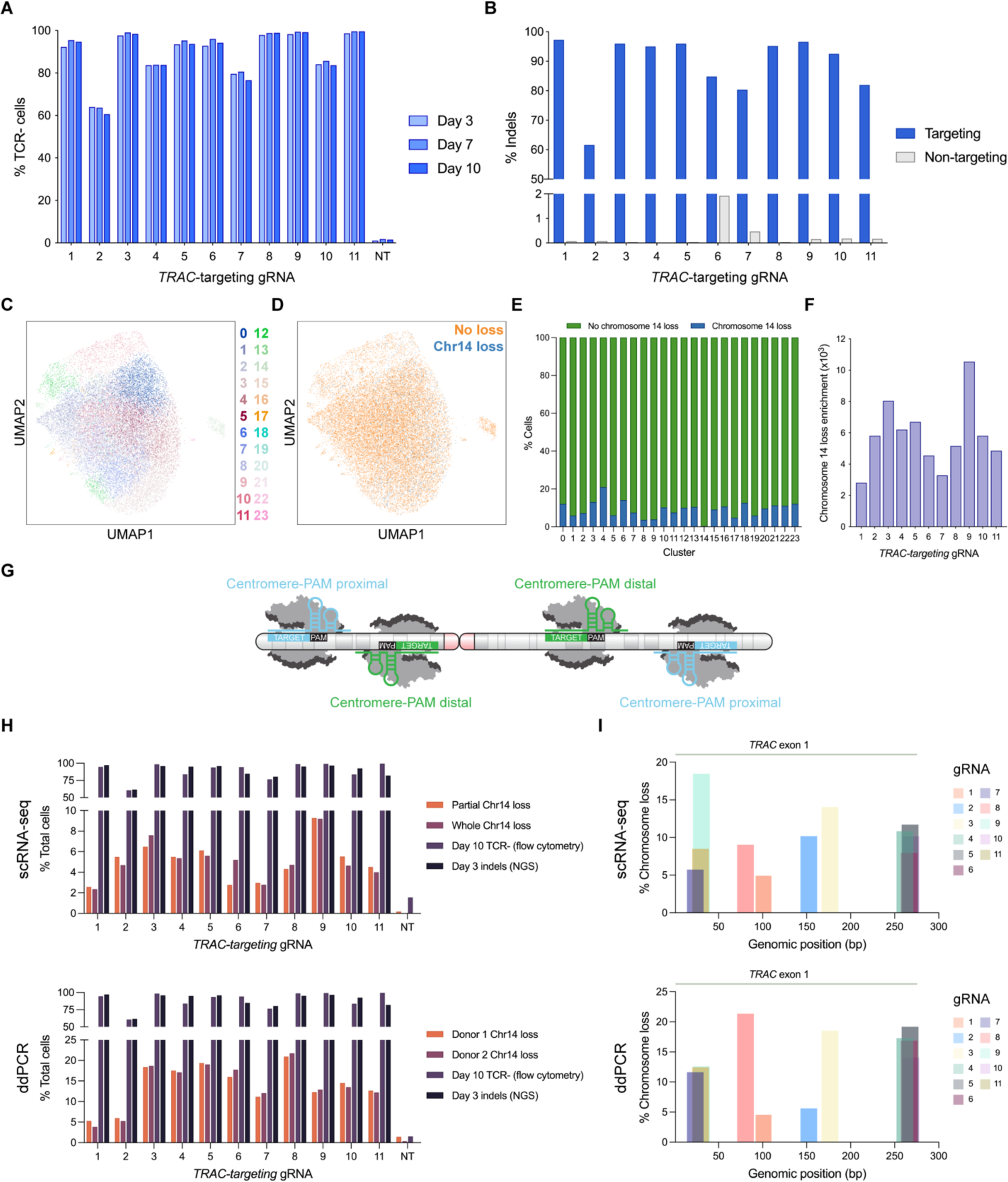
CRISPR-Cas9 genome editing results in chromosome loss regardless of guide RNA orientation or genomic position, related to Figure 1. (A) TCR disruption from Cas9 genome editing of *TRAC*. TCR expression was measured via flow cytometry at 3, 7, and 10 days post-nucleofection. NT indicates non-targeting gRNA. (B) Indels at the *TRAC* locus (targeting) from Cas9 genome editing as measured by next-generation sequencing. Cells treated with a non-targeting gRNA were evaluated for indels at each of the *TRAC* target sequences (non-targeting). (C) UMAP projection of T cells edited with a *TRAC*-targeting or non-targeting gRNA. (D) The UMAP projection of T cells within the *TRAC* editing experiment was overlayed with estimations of whether the cell had chromosome 14 loss or not (whole or partial chromosome loss). (E) Percentage of cells with chromosome 14 loss per cluster (see Fig. S1D). (F) Quantification of chromosome 14 loss enrichment across 11 different *TRAC*-targeting gRNAs. Chromosome 14 loss enrichment was calculated relative to T cells treated with Cas9 and a non-targeting gRNA. (G) Schematic of gRNA orientation relative to the centromere. Cas9 targets where the PAM was proximal to the centromere (red) relative to the target DNA sequence were considered centromere-PAM proximal (blue), while Cas9 targets where the PAM was distal to the centromere relative to the target DNA sequence were considered centromere-PAM distal (green). (H) Comparison of TCR disruption (by flow cytometry or next-generation sequencing) and chromosome 14 loss as measured by scRNA-seq (top) or ddPCR (bottom). (I) Chromosome 14 loss by scRNA-seq (combination of whole and partial chromosome 14 loss, top) or ddPCR (mean of n = 3 replicates, n = 2 biological donors, bottom) for each gRNA based on target position within the *TRAC* gene. Gray line indicates the first exon of *TRAC* (274 bp).

**Figure S2:**
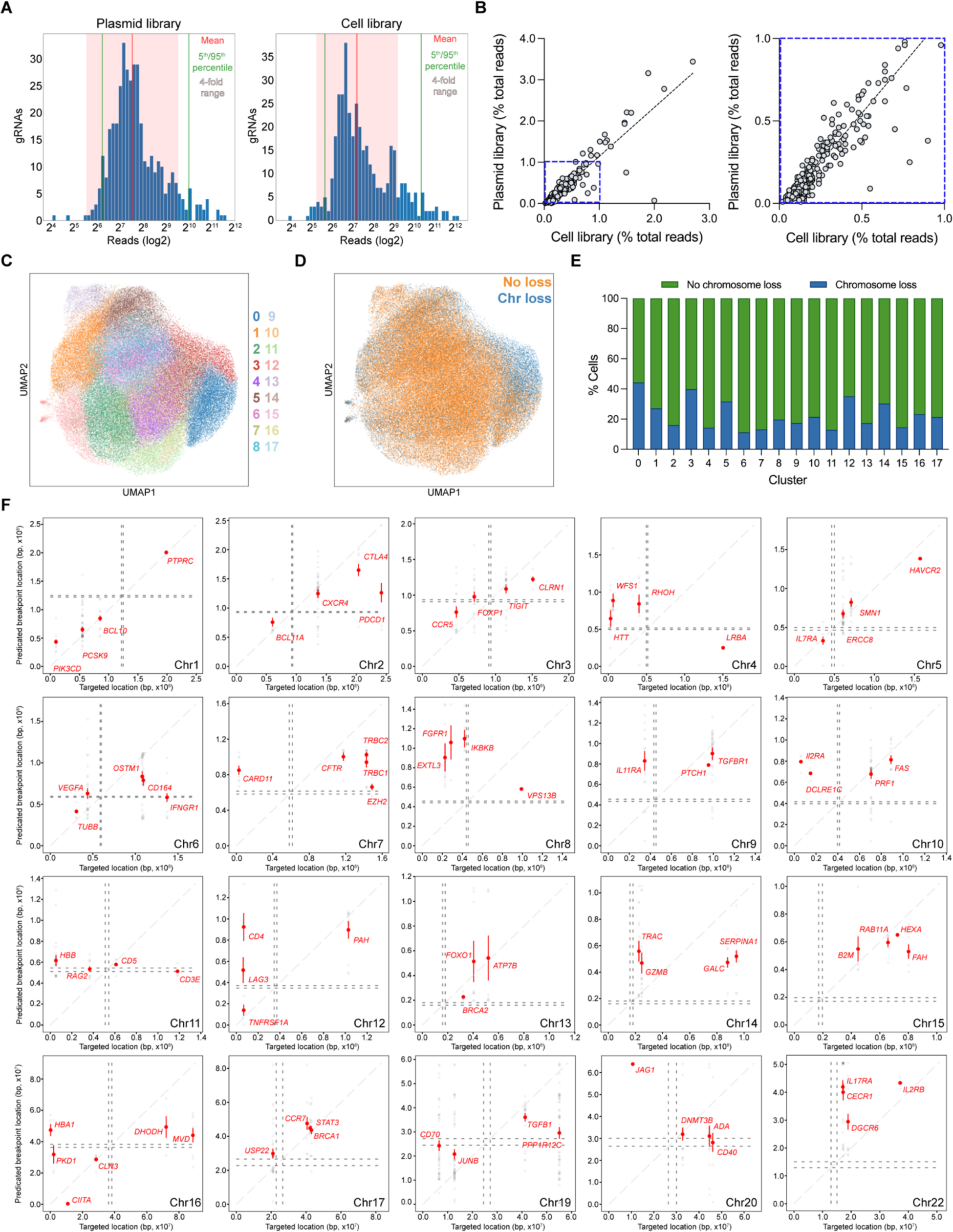
CROP-seq reveals genome-scale breakpoints and chromosome loss, related to Figure 2. **(A)** Distribution of next-generation sequencing reads for each gRNA in the CROP-seq gRNA library as a plasmid (left) or once integrated into T cells via lentivirus (right). The mean is shown as a red line, 5^th^ and 95^th^ quartiles are shown as green lines, and a 4-fold range to either side of the mean is shown as a pink shaded area. **(B)** Correlation between gRNA reads in the plasmid library and cell library. A zoomed in perspective (blue dashed line, left) is shown in a separate panel (blue dashed line, right). Dashed gray line represents the linear regression line of best fit (Slope = 1.189, R^2^ = 0.8348). **(C)** UMAP projection of T cells from the CROP-seq screen. **(D)** The UMAP projection of T cells within the CROP-seq screen was overlayed with estimations of whether the cell had a targeted chromosome loss or not (whole or partial chromosome loss). **(E)** Percentage of cells with targeted chromosome loss per cluster (see Fig. S2C). **(F)** Predicted breakpoint location versus intended gRNA target location from the CROP-seq screen. gRNAs are grouped by targeted chromosome. Chromosomes 18 and 21 are omitted because no partial chromosome loss was detected. Red data points represent the mean and 95% confidence interval.

**Figure S3:**
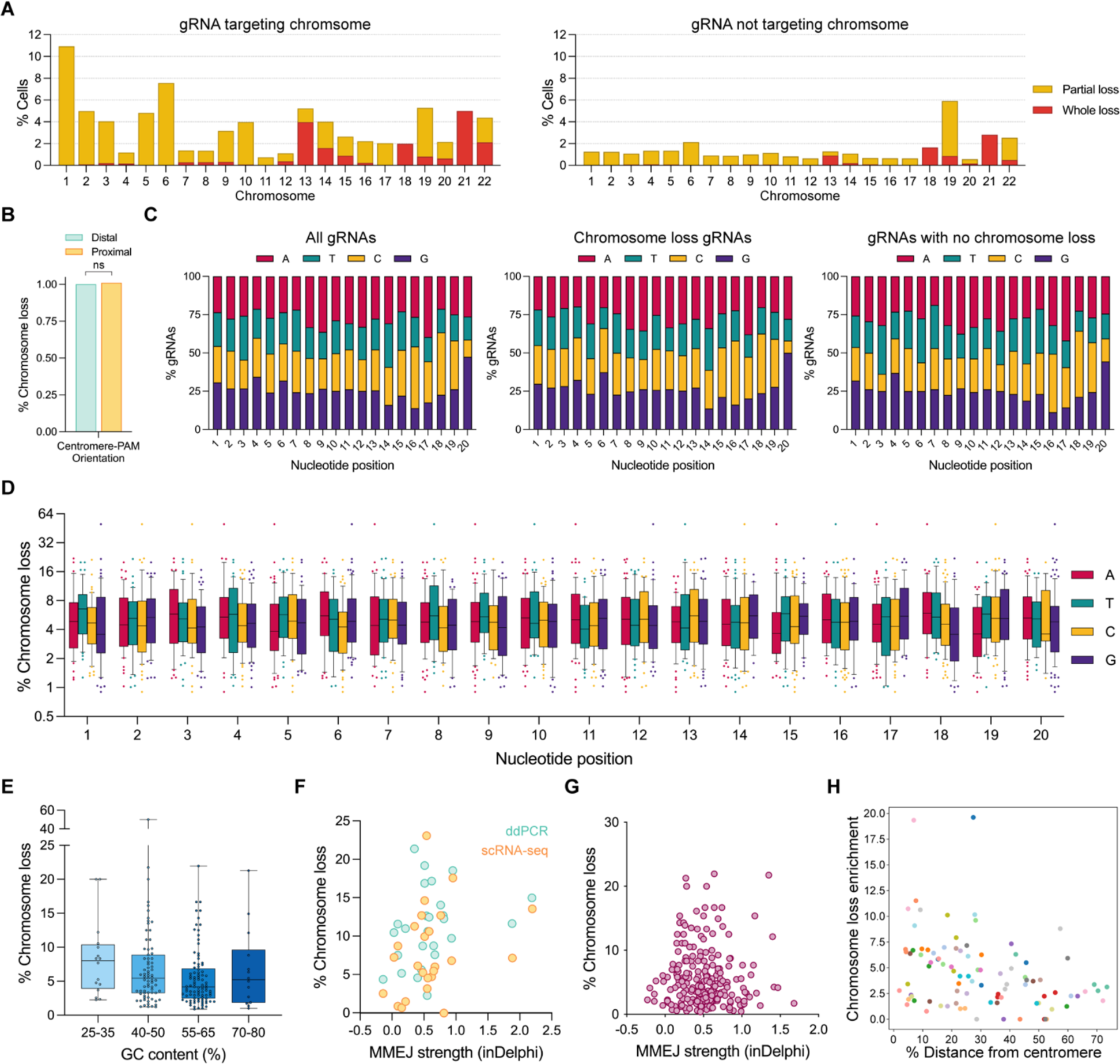
Influence of genetic context and Cas9 gRNA sequence on chromosome loss, related to Figure 2. **(A)** Partial and whole chromosome loss from the CROP-seq screen. Chromosome loss was quantified at chromosomes where the gRNA was targeting that specific chromosome (left) or at chromosomes not targeted by the gRNA (right). Chromosome loss at chromosomes not targeted by the gRNA was used to calculate baseline noise. **(B)** Influence of gRNA orientation on chromosome loss. gRNAs where the PAM is distal to the centromere relative to the gRNA target sequence were compared against gRNAs where the PAM is proximal to the centromere relative to the gRNA target sequence. *P*-value was calculated using a two-sided Fisher’s Exact Test and was 0.592413. ns = not significant. **(C)** Distribution of nucleotides across each position of the gRNA spacer within the CROP-seq library. Distribution for all gRNAs (left), gRNAs that resulted in chromosome loss (middle), and gRNAs that did not result in chromosome loss (right). **(D)** Chromosome loss by nucleotide identity across each position of the gRNA spacer within the library. **(E)** Chromosome loss by gRNA spacer sequence GC content. gRNAs were arbitrarily binned by varying levels of GC content. **(F)** Chromosome loss versus computationally predicted MMEJ influence for Cas9 RNP nucleofection experiments (teal = ddPCR measurements, yellow = scRNA-seq measurements. Chromosome loss rates are identical to Fig. 2e). ddPCR Spearman’s correlation = 0.40, \**P* = 0.04 (two-tailed); scRNA-seq Spearman’s correlation = 0.27, *P* = 0.19 (two-tailed). **(G)** Chromosome loss versus computationally predicted MMEJ influence for the CROP-seq screen experiment. gRNAs with non-zero chromosome loss were plotted. Spearman correlation = −0.08, *P* = 0.25 (two-tailed). **(H)** Chromosome loss by position along the target chromosome. Distance from the centromere was normalized to the length of the target chromosome. Spearman’s correlation = −0.34, \*\*\**P* = 0.0009 (two-tailed).

**Figure S4:**
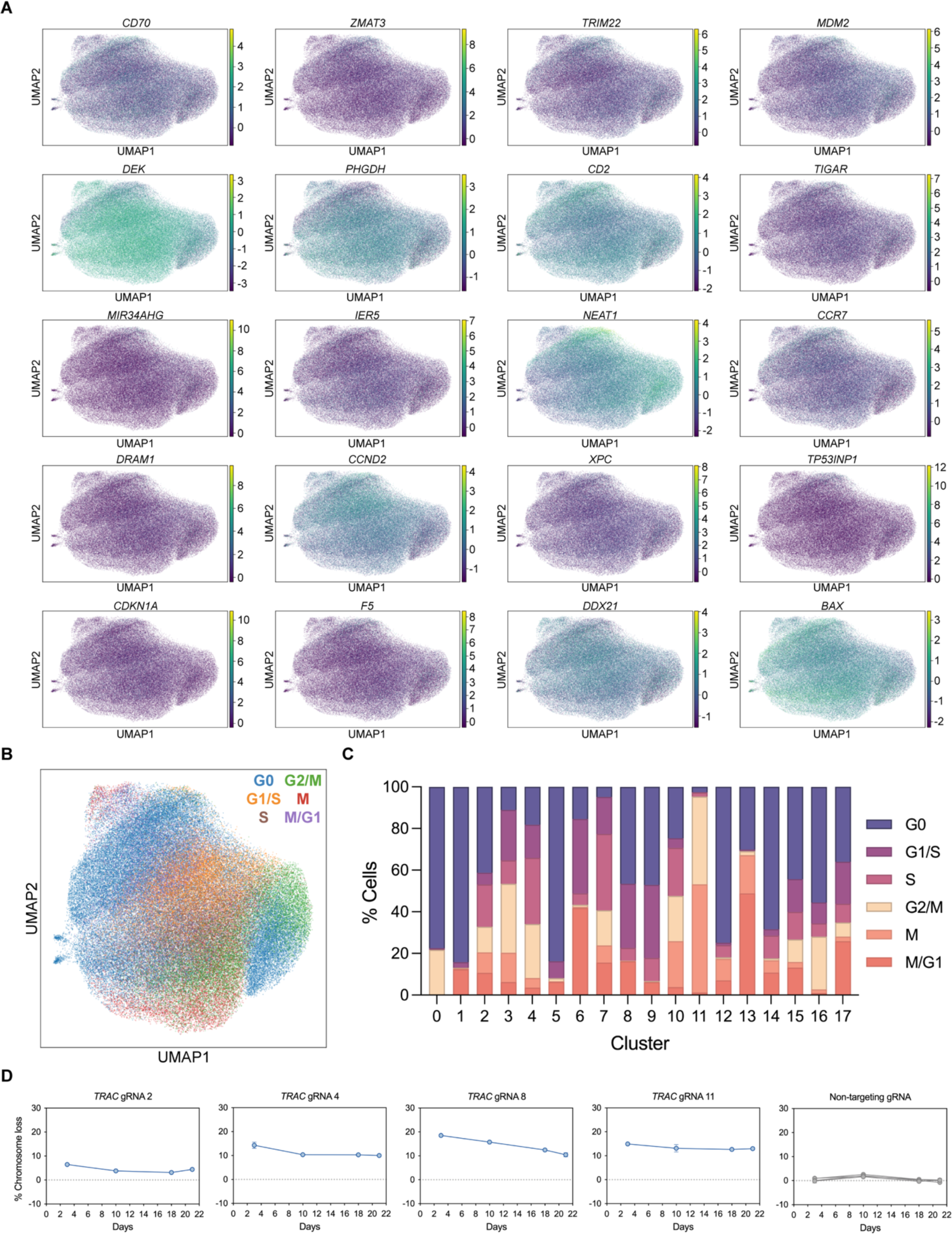
Cas9-induced chromosome loss is associated with differential gene expression and a fitness disadvantage, related to Figure 3 and Figure 4. **(A)** UMAP projections of T cells within the CROP-seq screen. Gene expression was overlayed onto the projections for the top 20 genes that were most differentially expressed across cells that had chromosome loss. **(B)** UMAP projection of T cells within the CROP-seq screen overlayed with cell cycle markers (see Fig. S2C). **(C)** Quantification of cell cycle states across the different clusters (see Fig. S2C). **(D)** ddPCR measurements of chromosome loss at the Cas9 *TRAC* target site over 21 days. Error bars represent the standard deviation from the mean (n = 3).

**Figure S5:**
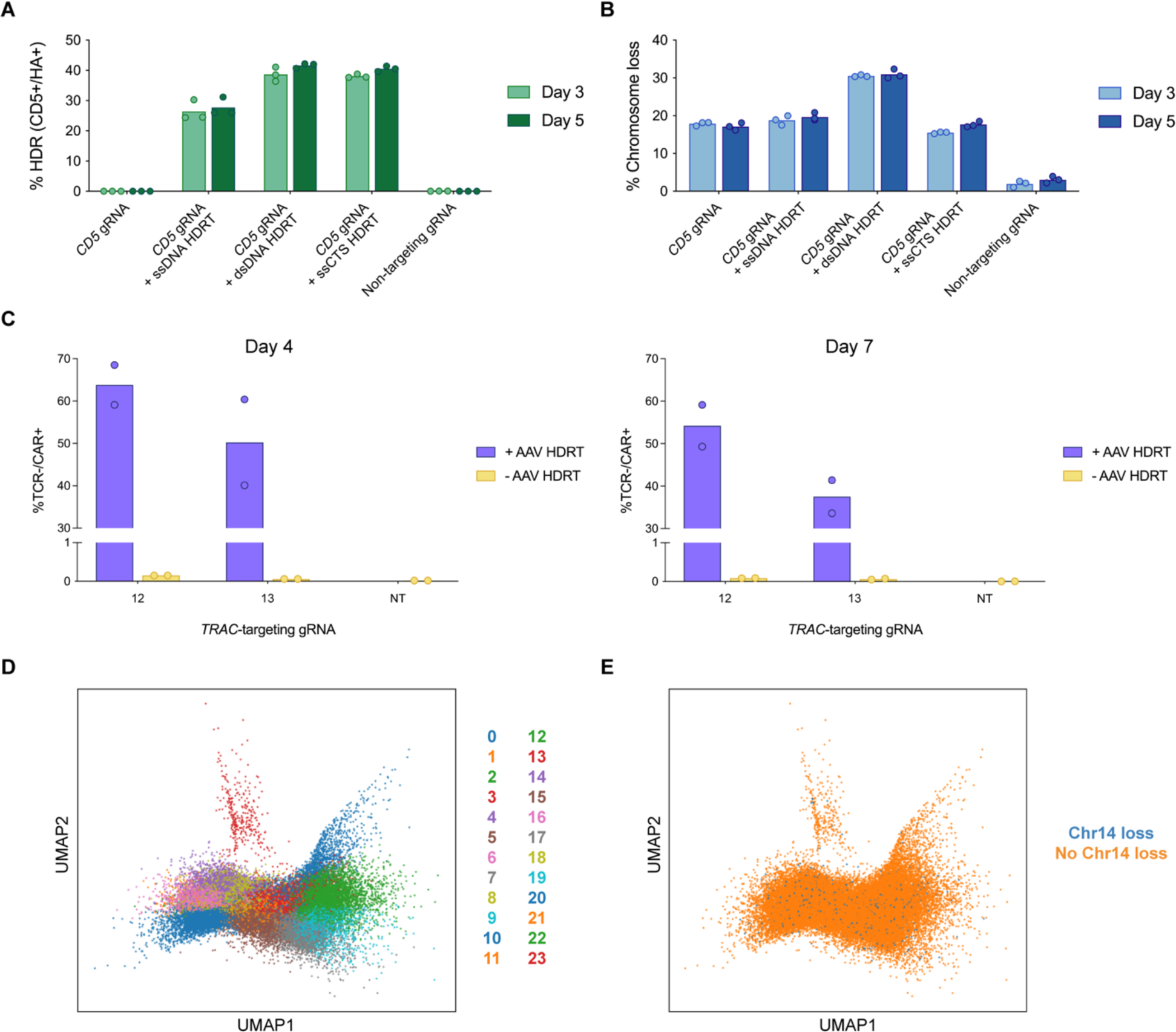
CRISPR-Cas9 homology-directed repair results in chromosome loss, related to Figure 5. **(A)** HDR efficacy 3 or 5 days post-nucleofection, determined by measuring CD5+/HA+ T cells via flow cytometry. **(B)** Chromosome loss at the target *CD5* locus via ddPCR, 3 or 5 days post-nucleofection. **(C)** HDR efficacy determined by measuring TCR-/CAR+ T cells via flow cytometry, four or seven days post-nucleofection. Two separate nucleofections/transductions were conducted for the different time points (n = 2 biological donors). **(D)** UMAP projection of CAR T cells generated via Cas9 HDR. Projection is an aggregate of two biological donors and multiple time points. **(E)** Distribution of CAR T cells with chromosome 14 loss across the UMAP projection (whole or partial chromosome loss).

**Figure S6:**
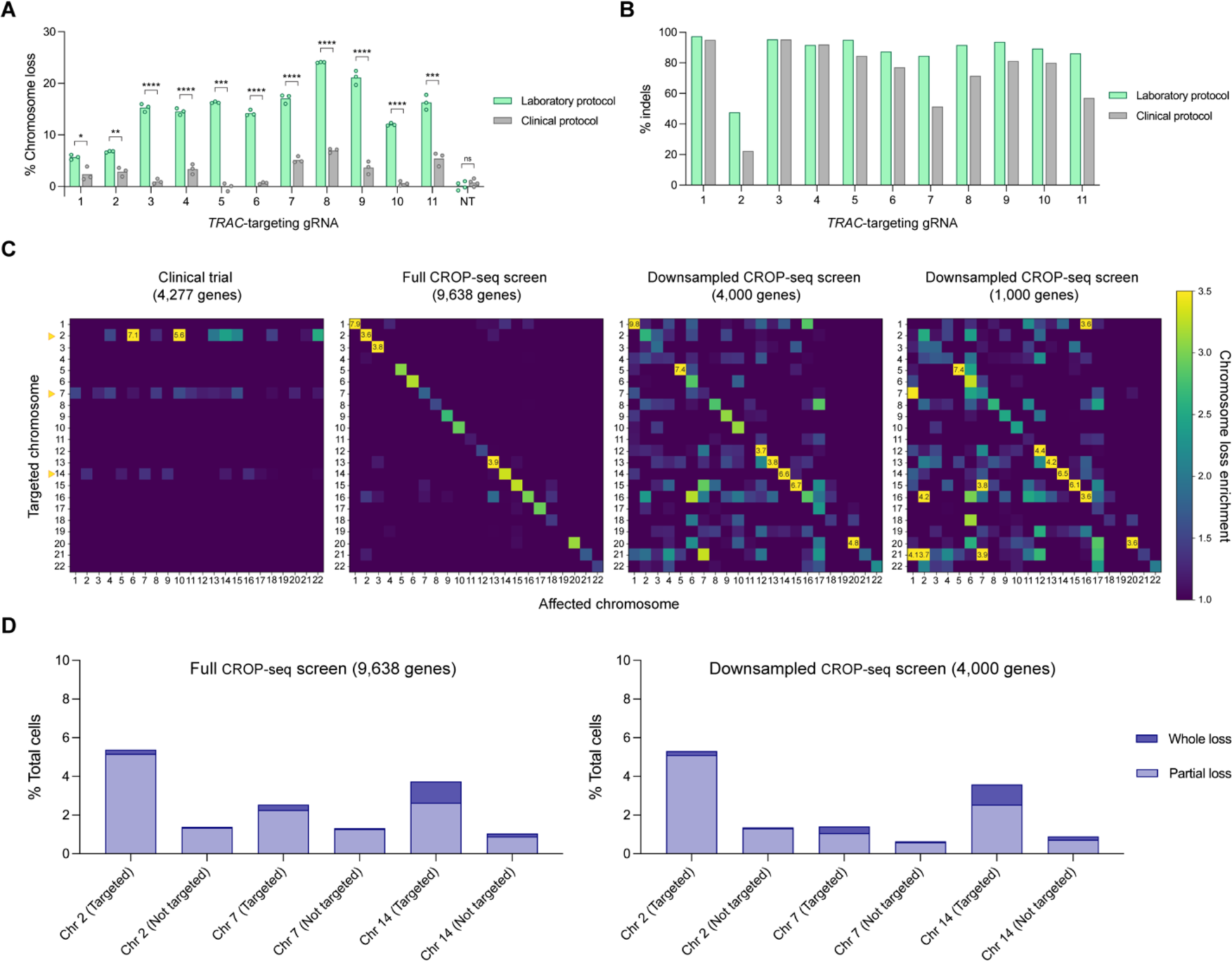
Clinical CRISPR-Cas9 genome editing protocol reduces chromosome loss in T cells, related to Figure 6. **(A)** Chromosome loss with 11 different *TRAC*-targeting gRNAs or a non-targeting gRNA (NT) using the laboratory or clinical protocol in T cells (n = 3). *P*-values are from Welch’s unpaired t-tests and from left to right are 0.010220, 0.004303, 0.000063, 0.000083, 0.000170, 0.000083, 0.000063, 0.000063, 0.000063, 0.000031, 0.000224, and 0.079286. **(B)** Indels measured by next-generation sequencing at the *TRAC* locus by Cas9 genome editing with the laboratory or clinical protocol. **(C)** Downsampling analysis to investigate the influence of total genes on chromosome loss enrichment. Rows represent the chromosome targeted by Cas9 and its gRNA. Columns represent the chromosome analyzed for chromosome loss. Chromosomal loss enrichment for all clinical trial patients and timepoints was evaluated only when targeting chromosomes 2 (*PDCD1*), 7 (*TRBC*), and 14 (*TRAC*) (yellow arrows, left heatmap). 4,277 total genes were detected in the clinical trial dataset. The full CROP-seq screen dataset (9,638 genes, second from left) was downsampled to 4,000 genes (second from right) or 1,000 genes (right) to evaluate the influence of total genes on chromosomal loss enrichment. **(D)** Quantification of whole and partial chromosome loss at chromosomes 2, 7, and 14 from the CROP-seq screen. Chromosome loss was measured at the specific chromosome targeted by the Cas9 gRNA (targeted) or at chromosomes not targeted by the Cas9 gRNA (not targeted), from the full CROP-seq screen dataset (left) or the CROP-seq screen dataset downsampled to mimic the clinical trial dataset (right).

